# Regulation of cyclic electron flow by chloroplast NADPH-dependent thioredoxin system

**DOI:** 10.1101/261560

**Authors:** Lauri Nikkanen, Jouni Toivola, Andrea Trotta, Manuel Guinea Diaz, Mikko Tikkanen, Eva-Mari Aro, Eevi Rintamäki

**Author notes:** The author responsible for distribution of materials integral to the findings presented in this article in accordance with the policy described in the Instructions for Authors (www.plantphysiol.org) is: Eevi Rintamäki. **Corresponding author:** Eevi Rintamäki Molecular Plant Biology Department of Biochemistry University of Turku FI-20014 TURKU Finland.

## Abstract

Linear electron transport in the thylakoid membrane drives both photosynthetic NADPH and ATP production, while cyclic electron flow (CEF) around photosystem I only promotes the translocation of protons from stroma to thylakoid lumen. The chloroplast NADH-dehydrogenase-like complex (NDH) participates in one CEF route transferring electrons from ferredoxin back to the plastoquinone pool with concomitant proton pumping to the lumen. CEF has been proposed to balance the ratio of ATP/NADPH production and to control the redox poise particularly in fluctuating light conditions, but the mechanisms regulating the NDH complex remain unknown. We have investigated potential regulation of the CEF pathways by the chloroplast NADPH-thioredoxin reductase (NTRC) *in vivo* by using an Arabidopsis knockout line of *NTRC* as well as lines overexpressing NTRC. Here we present biochemical and biophysical evidence showing that NTRC activates the NDH-dependent CEF and regulates the generation of proton motive force, thylakoid conductivity to protons and redox balance between the thylakoid electron transfer chain and the stroma during changes in light conditions. Further, protein–protein interaction assays suggest a putative thioredoxin-target site in close proximity to the ferredoxin binding domain of NDH, thus providing a plausible mechanism for regulation of the NDH ferredoxin:plastoquinone oxidoreductase activity by NTRC.

**One sentence summary:** Chloroplast thioredoxins regulate photosynthetic cyclic electron flow that balances the activities of light and carbon fixation reactions and improves plant fitness under fluctuating light conditions.

## INTRODUCTION

In their natural habitats, plants face constant fluctuation of light intensity, including both seasonal changes in photoperiod and daily fluctuations according to environmental conditions. Optimization of photosynthesis in plant leaves requires strict balancing between conversion of light energy to chemical energy in photosynthetic light reactions and the energy-consuming reactions of chloroplast metabolism. Multiple regulatory and photoprotective mechanisms have evolved in photosynthetic organisms to cope with fluctuating light conditions and to prevent the photodamage of both Photosystem (PS) II and PSI (Tikkanen et al., 2012; Tikkanen and Aro, 2014; Tiwari et al., 2016; Townsend et al., 2017). Regularly occurring light variations induce long-term acclimatory changes in the photosynthetic machinery via signaling mechanisms, while temporary fluctuation of light within a day transiently activates short-term regulatory mechanisms (Bailey et al., 2001; Grieco et al., 2012; Kono and Terashima, 2014; Armbruster et al., 2014). The short-term mechanisms include non-photochemical quenching (NPQ), photosynthetic control of electron flow between PSII and PSI, state transitions (ST), cyclic electron flow (CEF), and activation of photosynthetic enzymes both in light and carbon fixation reactions (Demmig-Adams et al., 2012; Tikkanen and Aro, 2014; Balsera et al., 2014; Yamori et al., 2016; Gollan et al., 2017).

Light drives the electron flow from water through PSII, plastoquinone (PQ), cytochrome *b6f*, plastocyanin (PC) and PSI to ferredoxin and ultimately to NADP^+^, producing NADPH. These photosynthetic electron transfer reactions are coupled to ATP synthesis via translocation of protons to the thylakoid lumen, generating a proton gradient over the thylakoid membrane (ΔpH), which together with membrane potential (ΔΨ) constitutes the proton motive force (*pmf*) (Hangarter and Good, 1982; Armbruster et al., 2017). ΔpH also contributes to induction of the energy-dependent qE component of NPQ, a photoprotective mechanism that dissipates excess excitation energy from the electron transfer chain (Niyogi and Truong, 2013; Ruban, 2016), and maintains photosynthetic control at Cyt *b6f* (Joliot and Johnson, 2011; Johnson, 2011). Other regulatory mechanisms include the reversible rearrangements of light harvesting complexes to balance the excitation of PSII and PSI known as state transitions (Tikkanen et al., 2006; Ruban and Johnson, 2009; Rochaix, 2011) as well as cyclic electron flow around PSI (CEF), a process where electrons are transferred from ferredoxin back to the PQ pool. CEF contributes to generation of *pmf* and therefore to production of ATP, and has been suggested to adjust the ATP/NADPH ratio in chloroplasts according to the needs of the CBC (for a recent review, see Yamori and Shikanai (2016). CEF provides an alternative electron acceptor mechanism for PSI to relieve stromal over-reduction, which is needed to protect the photosystems from damage during early developmental stages of chloroplasts (Allorent et al., 2015; Suorsa, 2015), and during excess illumination or fluctuating light conditions (Miyake et al., 2004; Suorsa et al., 2012; Yamori and Shikanai, 2016; Yamori et al., 2016). CEF has also been shown to be important for controlling the magnitude of the *pmf* (Wang et al., 2015; Shikanai and Yamamoto, 2017), and during induction of photosynthesis (Joliot and Joliot, 2002; Fan et al., 2007). Fan et al. (2007) calculated that CEF contributes a maximum of 68% of total electron flux after 30 s illumination of spinach leaves with red and far red light.

Two distinct pathways of CEF have been suggested to exist in plant chloroplasts (Munekage et al., 2004). One CEF pathway involves the chloroplast NADH dehydrogenase-like complex (NDH), an orthologue of mitochondrial respiratory complex I (Shikanai, 2016; Peltier et al., 2016). However, unlike complex I, which is reduced by NADH, the chloroplast NDH complex is reduced by ferredoxin (Yamamoto et al., 2011; Yamamoto and Shikanai, 2013). It has been suggested recently in several studies that CEF via the NDH complex is essential for photosynthesis in low light conditions (Yamori et al., 2015; Kou et al., 2015; Martin et al., 2015) as well as for the tolerance of drought (Horvath et al., 2000) and low temperature (Yamori et al., 2011). The antimycin A –sensitive CEF pathway depends on the proteins PROTON GRADIENT REGULATION 5 (PGR5) (Munekage et al., 2002) and PGR5-LIKE 1 (PGRL1) (DalCorso et al., 2008), and has been suggested to constitute the hypothetical ferredoxin-plastoquinone reductase (FQR) (Hertle et al., 2013). However, controversy still exists over the molecular identity of FQR and the physiological function of PGR5 (Leister and Shikanai, 2013; Tikkanen and Aro, 2014; Kanazawa et al. 2017). The PGR- and NDH-dependent pathways differ in their energetic properties; two protons per electron are translocated to the lumen (by the Q-cycle) in the FQR-pathway, whereas the NDH-complex functions as a proton pump and additionally transfers 2H+ per electron to the lumen (Strand et al., 2017). A third CEF pathway involving transfer of electrons from ferredoxin or FNR to PQ via heme c_n_ in the Cyt *b_6_f* complex has also been proposed (Hasan et al., 2013). In general, CEF activity is highly dependent on stromal redox state (Breyton et al., 2006), and both the PGR-dependent pathway (Hertle et al., 2013; Strand et al., 2016a) and the NDH pathway (Courteille et al., 2013) have been proposed to be subject to thiol-regulation by chloroplast thioredoxins.

In chloroplasts of Arabidopsis, two thioredoxin systems function in parallel. The Ferredoxin-thioredoxin system depends on photosynthetically reduced ferredoxin to supply electrons to the Ferredoxin-thioredoxin reductase (FTR), which in turn reduces several thioredoxins, namely TRX-*f*1 and *f*2, four isoforms of TRX-*m*, TRX-*x* as well as TRX-*y*1 and *y*2 (Schürmann and Buchanan, 2008; Yoshida and Hisabori, 2017). The other system consists of a single enzyme, NADPH-thioredoxin reductase (NTRC) that contains both a reductase and a thioredoxin domain (Serrato et al., 2004). NTRC is reduced by NADPH, which is produced, besides in the light reactions, also in the oxidative pentose phosphate pathway (OPPP) in darkness. Both chloroplast TRX systems are essential for normal development and growth of plants (Serrato et al., 2004; Wang et al., 2014). The *ntrc* knockout has a stunted and low chlorophyll phenotype, which is particularly severe in plants grown under short photoperiods (Perez-Ruiz et al., 2006; Lepistö et al., 2009; Lepistö et al., 2013). The mutant suffers from impaired ability to activate the ATP synthase and CBC enzymes as well as elevated nonphotochemical quenching (NPQ) (Nikkanen et al., 2016; Carrillo et al., 2016; Naranjo et al., 2016; Thormählen et al., 2017). In contrast, NTRC overexpression lines (OE-NTRC), with 15–20 times higher NTRC content compared to WT, show enhanced vegetative growth and increased activation of the ATP synthase and CBC enzymes, particularly in darkness and low light (Toivola et al., 2013; Nikkanen et al., 2016). NTRC has a less negative midpoint redox potential than FTR (Hirasawa et al., 1999; Yoshida and Hisabori, 2016) and plays an important regulatory role under low irradiance, while the FTR-dependent system probably requires more extensive illumination to be fully activated (Thormählen et al., 2017; Geigenberger et al., 2017; Nikkanen et al., 2016). Recent studies have revealed significant functional overlap and crosstalk between the two chloroplast TRX systems, and indicated that they cooperatively regulate ATP synthesis, the CBC, starch synthesis and scavenging of reactive oxygen species (ROS) (Thormählen et al., 2015; Nikkanen et al., 2016; Pérez-Ruiz et al., 2017; Geigenberger et al., 2017). Moreover, redox-regulation of both CEF pathways has been previously reported (Courteille et al., 2013; Hertle et al., 2013; Strand et al., 2016a). The physiological roles of each CEF pathway and TRXs involved in the regulation are nevertheless still unclear.

Here we have used the *ntrc* knockout mutant as well as NTRC overexpression lines of *Arabidopsis thaliana* to investigate the potential role of the NTRC system in regulating CEF. Our results emphasize the important role of thioredoxins in the chloroplast regulatory network, particularly controlling the photosynthetic redox balance under fluctuating light conditions. NTRC plays a crucial role in activation of the NDH-dependent CEF in darkness (chlororespiration) and during dark to light transitions. Overexpression of NTRC, on the other hand, maintains constant NDH-CEF activity leading to elevated *pmf* and improved utilization of light energy under fluctuating light conditions. Our results also suggest that NTRC does not activate the PGR-dependent CEF, but contributes to the PGR5-dependent downregulation of thylakoid membrane proton conductivity upon transient exposure of leaves to high light intensity. Through control of both CEF and the activity of the ATP synthase, NTRC plays a pivotal role in adjusting the proton motive force and photosynthetic redox poise in Arabidopsis chloroplasts.

## RESULTS

### NTRC is an active reductant in darkness and low light conditions

NADPH produced in the oxidative pentose phosphate pathway (OPPP) has been proposed to maintain the NTRC pool partially reduced, and thus active in darkness and when low irradiance limits photosynthesis (Perez-Ruiz et al., 2006; Geigenberger et al., 2017). To confirm this hypothesis, we analyzed the *in vivo* redox state of NTRC by a mobility shift assay using the WT or OE-NTRC protein extracts alkylated with methoxypolyethylene glycol maleimide (MAL-PEG). The assays indicated that the redox state of the NTRC pool remains fairly constant in all light intensities and during dark-to-light transitions, with a significant proportion of the enzyme pool in fully or partially reduced form (Fig. 1). This is also the case in OE-NTRC, despite the increase in NTRC content of leaves (Fig. 1, Suppl. Fig. S1). These results are in agreement with the hypothesis that NTRC acts as a thiol regulator of photosynthesis and chloroplast metabolism in darkness and low light conditions (Nikkanen et al., 2016; Carrillo et al., 2016; Thormählen et al., 2017).

**Figure 1.**
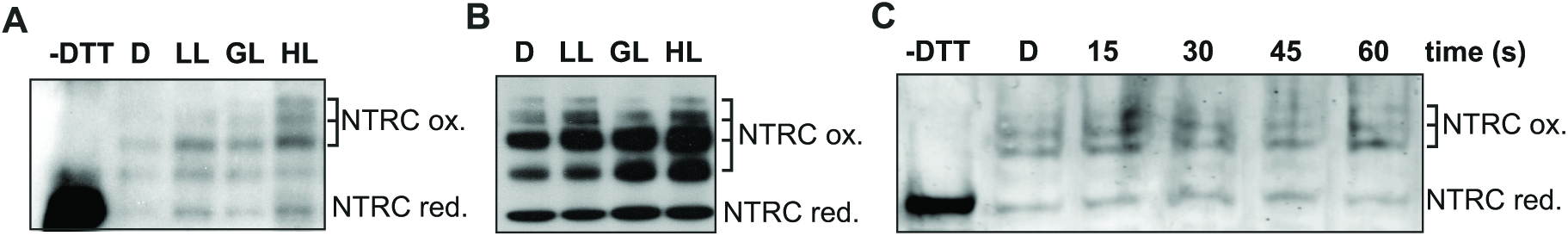
*In vivo* redox state of NTRC in dark-adapted and illuminated leaves. (**A**) and (**B**) Total protein extract was isolated from WT (A) and OE-NTRC (B) leaves incubated in darkness (D), or illuminated for 2h in low light (LL, 40 μmol photons m^−2^s^−1^), growth light (GL, 200 μmol photons m^−2^s^−1^) or high light (HL, 800 μmol photons m^−2^s^−1^). Free thiols of proteins were blocked with NEM, disulfides reduced with DTT and newly formed thiols alkylated with MAL-PEG. The in vivo-reduced form of NTRC therefore migrates faster in SDS-PAGE than the in vivo oxidized forms. –DTT stands for the unlabeled control sample where DTT was not added after incubating the leaf extracts in a buffer containing NEM. Protein content of samples has been equalized only based on the amount of starting leaf material, and the apparent differences in band intensity should not be taken as indication of differences in NTRC content between light treatments. For an analysis of the origin of different MAL-PEG labelled bands see Suppl. Fig. S1. (**C**) NTRC redox state in WT during a transition from dark to growth light. Samples were taken from darkness (2h) (D) and 15, 30, 45 and 60 seconds after onset of illumination.

### NDH-dependent CEF is enhanced by overexpression of NTRC

In order to determine the effect of altered chloroplast thiol-redox state on the activity of NDH-dependent CEF, we measured the post-illumination rise of chlorophyll a fluorescence (PIFR). The PIFR has been suggested to represent electron flow from stromal reductants via the NDH-complex to the plastoquinone (PQ) pool upon cessation of illumination (Shikanai et al., 1998; Gotoh et al., 2010). The OE-NTRC line showed a significantly larger PIFR after pre-illumination with low intensity white light than WT, suggesting increased CEF activity (Fig. 2). In agreement with previous reports, no PIFR was detected in the *ndho* mutant, which is lacking a functional NDH complex (Rumeau et al., 2005), while a diminished PIFR was observed in the *pgr5* line, which is deficient in PGR-dependent CEF (Munekage et al., 2002) (Fig. 2). These results suggest that NTRC contributes to activation of NDH-dependent CEF. In order to confirm that the increased PIFR in OE-NTRC derives from the activity of the NDH complex, we generated an NTRC overexpression line in the *ndho* mutant background (OENTRC *ndho*), which indeed was fully missing the PIFR (Fig. 2). The level of NTRC overexpression in OE-NTRC *ndho* plants was confirmed by immunoblotting and found to be similar to the OE-NTRC line (Suppl. Fig. S1).

**Figure 2.**
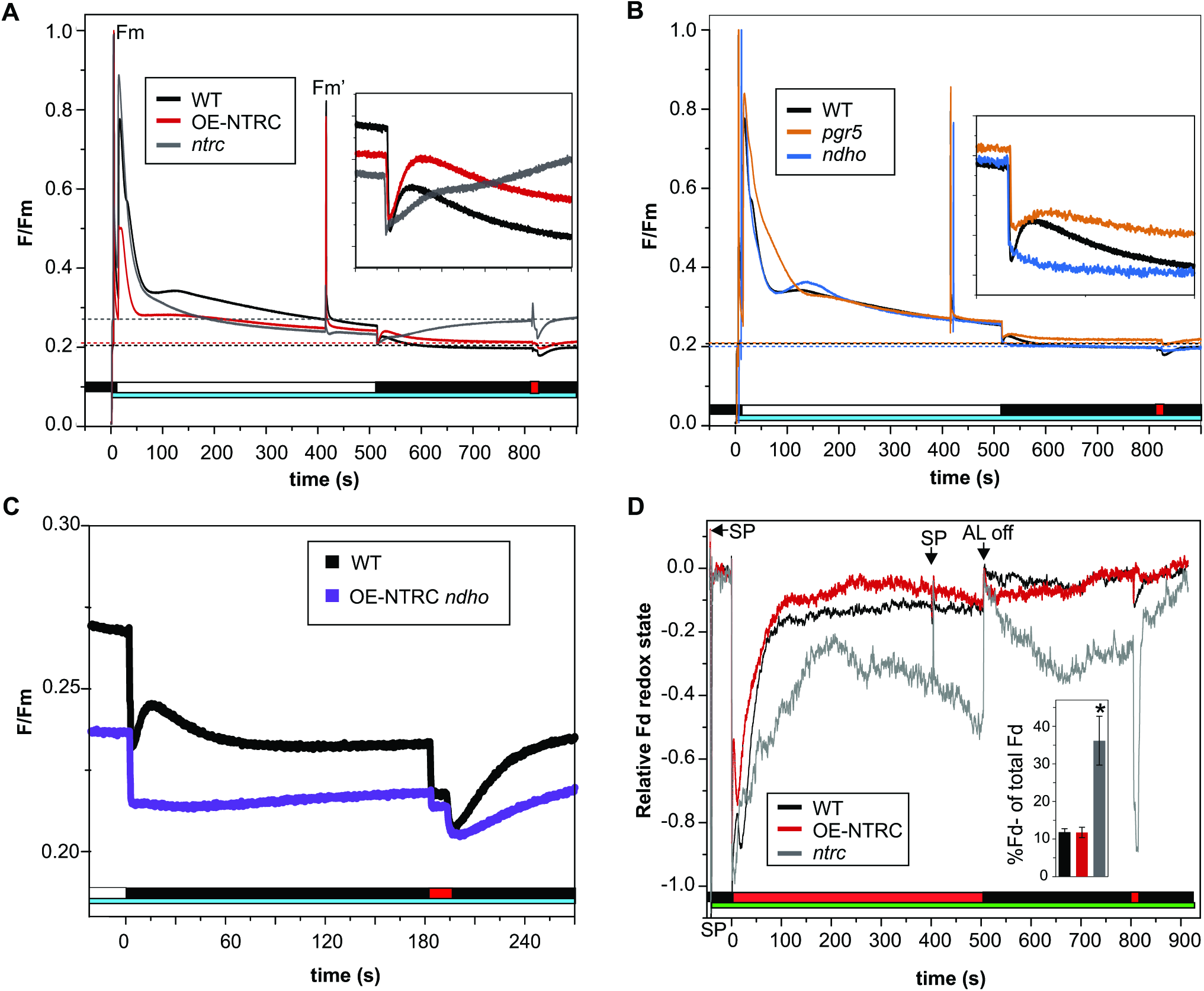
Post-illumination fluorescence rise (PIFR) in dark-adapted leaves. **(A)** and **(B)** PIFR was measured from WT, OE-NTRC, *ntrc* (A), *pgr5* and *ndho* (B) leaves. The smaller windows show magnifications of the ~100 s of the PIFR. The cyan bars indicate exposure to a 480 nm measuring light of 0.28 μmol photons m^−2^s^−1^, the white bar depicts illumination with 67 μmol photons m^−2^s^−1^ white light and the red bar shows the duration of a pulse of far red light. The dashed lines indicate the F_0_ values of the lines. The curves are averages of measurements from 3 to 7 individual leaves. (**C**) PIFR in WT and OE-NTRC *ndho*. Only the post-illumination phase of the experiment is shown in the figure. The curves are averages of measurements from 3 to 4 individual leaves. **(D)** Fd redox changes during and after illumination of dark-adapted WT, *ntrc* and OE-NTRC leaves. The Fd redox changes were deconvoluted from four near-infrared absorbance differences measured with a Dual/Klas-NIR spectrometer according to Klughammer and Schreiber (2016). Leaves were illuminated at actinic red light (630 nm) of 61 μmol photons m^−2^s^−1^. The red bar shows the duration of a pulse of far red light, the green bar the duration of the four measuring beams, the black bar the duration of the dark period after illumination, SP means saturating pulse. Averages of the relative amount of reduced Fd at the end of actinic illumination ± SE in 3 to 7 individual leaves are shown in the column chart inside the figure. Representative curves of WT, OE-NTRC and *ntrc* are shown in the figure.

The *ntrc* knockout exhibited a slower initial PIFR response after 500 s of light, but the PIFR did not decline after 15–20 s as in WT or OE-NTRC, but instead continued to rise throughout the duration of the dark phase of the experiment (Fig. 2). A brief pulse of far red (FR) light quenched the fluorescence, but after cessation of the FR light, fluorescence quickly rose back to its high pre-FR level. The F_0_ level was elevated in dark-adapted *ntrc* leaves compared to WT, OE-NTRC or other mutant lines (Fig. 2). The abnormal fluorescence pattern in Fig. 2 may also be due to the highly pleiotropic phenotype of the *ntrc* mutant, particularly when grown under a short day photoperiod. The *ntrc* knockout had high NPQ in the experimental conditions (Suppl. Fig. S2), and its relaxation in darkness likely contributed to the PIFR. Moreover, the *ntrc* mutant has an impaired capacity to scavenge hydrogen peroxide (H_2_O_2_) (Kirchsteiger et al., 2009, Pulido et al., 2010), whose accumulation has been shown to cause an increase in NDH-dependent CEF (Strand et al., 2015, Strand et al., 2017b). In order to clarify whether the differences in PIFR were caused indirectly by metabolic disturbances due to impaired growth in a short photoperiod and/or accumulation of H_2_O_2_, we estimated the levels of H_2_O_2_ in illuminated WT, *ntrc* and OE-NTRC leaves by DAB staining, and repeated the PIFR-experiment with plants grown in a 12h/12h photoperiod. An increased amount of H_2_O_2_ was detected in both low light and high light-treated *ntrc* leaves in comparison to WT (Suppl. Fig. S2), in agreement with Pulido et al. (2010). No difference was observed in the amount of H_2_O_2_ between OE-NTRC and WT (Suppl. Fig. S2), indicating that the elevation of PIFR in OE-NTRC (Fig. 2) is not caused by increased content of H_2_O_2_. Furthermore, the PIFR mostly disappeared in 12h photoperiod grown *ntrc*, but remained similar to 8h photoperiod grown plants in WT and OE-NTRC (Suppl. Fig. S2).

Altered PIFR responses in OE-NTRC and *ntrc* could hypothetically be caused by changes in the available amount of reduced ferredoxin (Fd^−^), the substrate of the NDH-complex (Yamamoto et al., 2011; Yamamoto and Shikanai, 2013). In order to investigate this possibility, we used the Dual/Klas-NIR spectrometer, which allows deconvolution of the Fd signal from plastocyanin (PC) and P700 signals (Klughammer and Schreiber, 2016; Schreiber and Klughammer, 2016; Schreiber, 2017), to measure the redox state of Fd in WT, *ntrc* and OE-NTRC under similar light- and post-illumination conditions as used in the PIFR measurements. In OE-NTRC, reduced Fd was re-oxidized slightly faster than in WT upon onset of illumination, but upon cessation of actinic illumination, there was no difference in the fraction of reduced Fd between WT and OE-NTRC (~11 % in both lines, Fig. 2). In *ntrc* leaves, however, almost 40 % of the Fd pool was reduced at the end of actinic illumination, and significantly increased reduction of Fd also occurred in darkness after initial oxidation (Fig. 2). These results indicate that the elevated PIFR in OE-NTRC is not caused by increased accumulation of substrate of the NDH complex. In contrast, the slow-rising high PIFR in *ntrc* is very likely partially caused by an increase in the relative amount of reduced Fd at the end of illumination.

Differences in the rate of PQ reduction could also be caused by altered content of PSII or PSI complexes, the NDH complex, Cyt *b6f* or plastid terminal oxidase (PTOX). No statistically significant differences were detected in the amounts of the PSII core protein D1, Cyt *b6f* subunit Cyt *f* or NDH subunits NhdS and NdhH between the studied lines, while a decrease in the amount of PGR5 and the PSI core protein PsaB, and an elevated PTOX content were detected in *ntrc* (Fig. 3).

**Figure 3.**
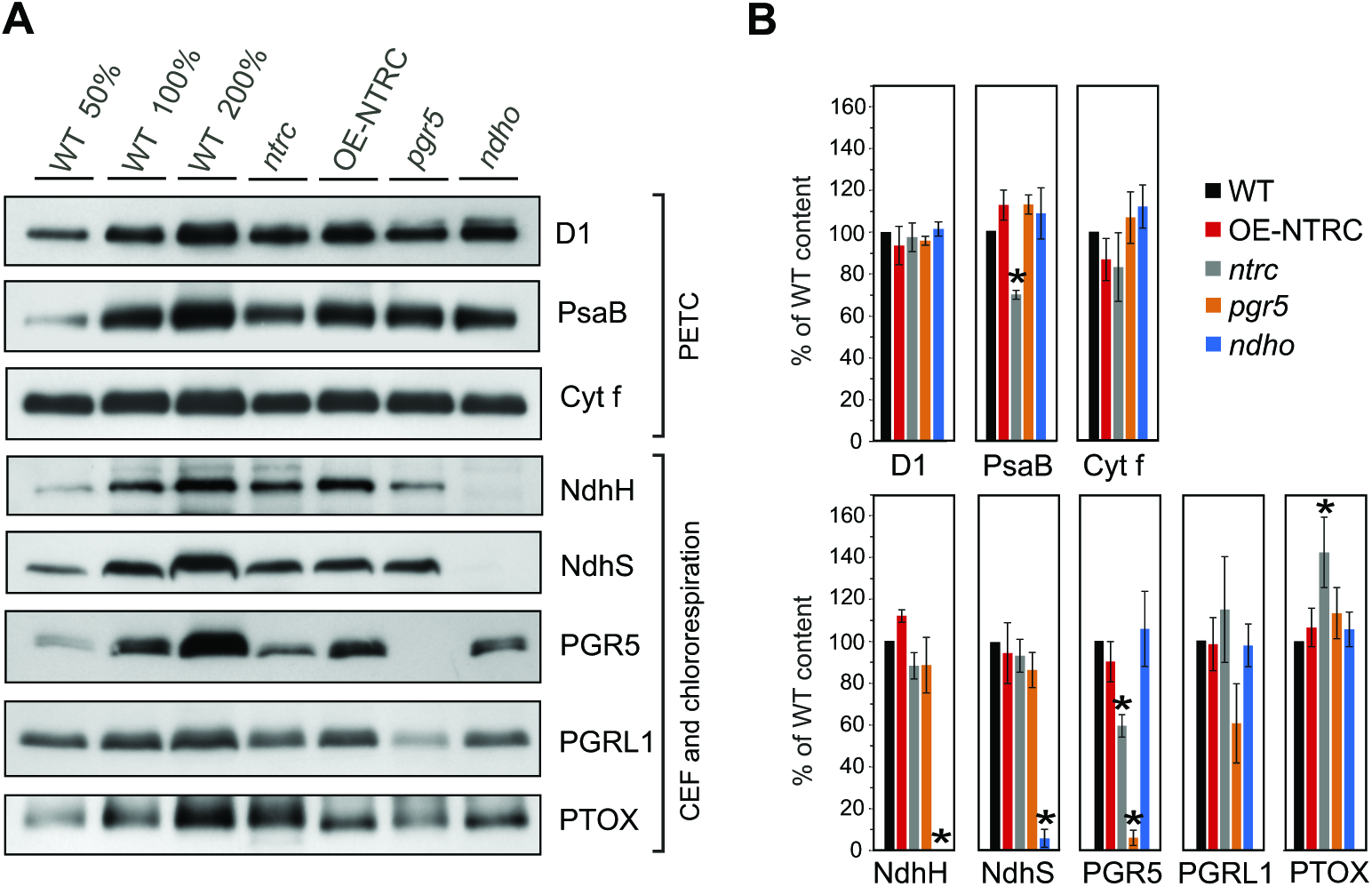
Content of proteins functioning in the photosynthetic electron transfer chain (PETC), cyclic electron flow (CEF) and chlororespiration in WT, *ntrc*, OE-NTRC, *pgr5* and *ndho*. **(A)** Representative immunoblots showing the content of D1, PsaB, Cyt *f*, NdhH, NdhS, PGR5, PGRL1 and PTOX. Appropriate amount of thylakoid extract (based on protein content) was separated with SDS-PAGE and probed with specific antibodies. Equal loading was confirmed by protein staining with Li-Cor Revert Total Protein Stain. **(B)** Relative content of proteins in mutant lines as percentage of WT. The numbers represent the average protein content ±SE in 3 to 5 biological replicates. The quantified values were normalized to the total protein content in the sample determined with Li-Cor Revert Total Protein Stain. Statistically significant differences to WT according to Student’s T-tests (P<0.05) are marked with *.

### NTRC promotes dark-reduction of the plastoquinone pool

To determine whether the higher CEF activity, observed in darkness after illumination of OE-NTRC (Fig. 2) alters the redox state of the PQ pool, we proceeded to analyze the phosphorylation level of LHCII proteins in dark-adapted and illuminated leaves to determine if the higher CEF activity alters the redox state of the PQ pool. Reduction of the PQ pool induces phosphorylation of LHCII by activating the STN7 kinase through interaction with the Cyt *b6f* complex (Vener et al., 1997; Bellafiore et al., 2005; Shapiguzov et al., 2016). In WT LHCII proteins were mostly non-phosphorylated in darkness, maximally phosphorylated in low light, moderately phosphorylated in growth light and mostly dephosphorylated in high light (Fig. 4H), in agreement with earlier studies (Rintamäki et al., 1997; Tikkanen et al., 2010). The *stn7* mutant was unable to phosphorylate LHCII (Bellafiore et al., 2005). In contrast to WT, LHCII was phosphorylated in darkness in OE-NTRC (Fig. 4H). As in WT, only a small amount of phosphorylated LHCII was present in thylakoids isolated from dark-adapted leaves of *ntrc*. Interestingly, de-phosphorylation of LHCII proteins in high light was slightly impaired in *ntrc* (Fig. 4). This is in accordance with earlier studies showing that phosphorylation of LHCII is highly dependent on stromal thiol redox state in high light (Rintamäki et al., 2000; Martinsuo et al., 2003).

**Figure 4.**
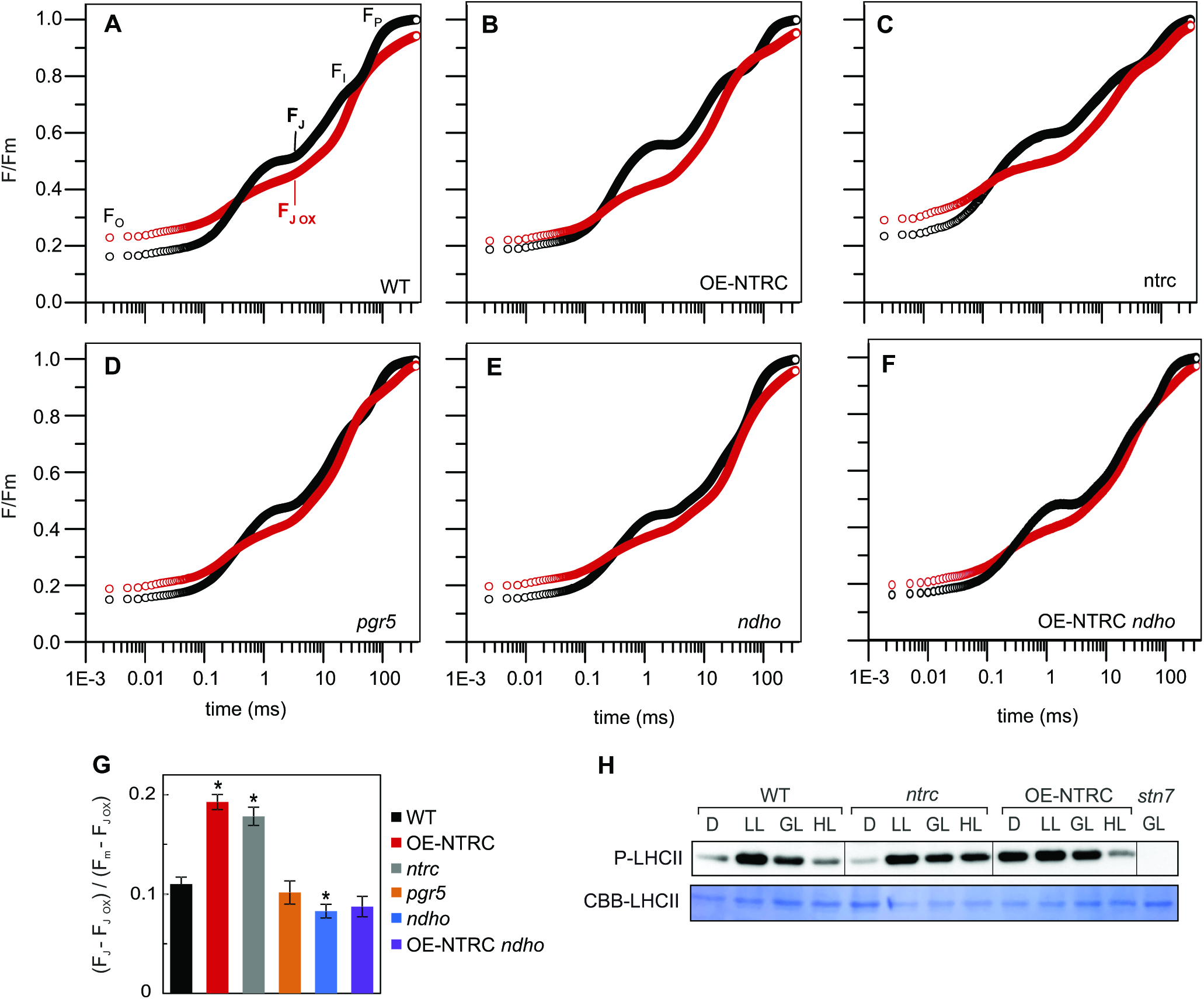
Redox state of the PQ pool in dark-adapted leaves. **(A–F)** PQ pool redox state in darkness was determined from measurement of the OJIP transients in dark-adapted leaves (black) and leaves pre-illuminated with far red light (red). Data is presented as averaged curves from measurements of 5 to 10 individual leaves on a logarithmic time scale from WT (A), OE-NTRC (B), *ntrc* (C), *pgr5* (D), *ndho* (E) and OE-NTRC *ndho* (F). The slight rise in F0 values after FR-illumination is likely due to the small actinic effect of FR on PSII, as discussed by Schansker and Strasser (2005). **(G)** Proportions of reduced Q_A_ calculated as (F_J_-F_J ox_)/ (F_m_-F_J ox_). Values are averages of measurements from 5 to 10 individual leaves ± SE. * indicates statistically significant difference to WT according to Student’s T test (P<0.05). All values are normalized to dark-adapted Fm. **(H)** Determination of phosphorylation status of LHCII proteins in WT, *ntrc*, OE-NTRC and *stn7* after 2 h of darkness and in low light (40 μmol photons m^−2^s^−1^), growth light (200 μmol photons m^−2^s^−1^) and high light (600 μmol photons m^−2^s^−1^). Thylakoid extracts containing 0.4 μg chlorophyll were separated with SDS-PAGE and detected with a Phosphothreonine-specific antibody. Coomassie Brilliant Blue staining of LHCII on the membrane (CBB-LHCII) was used as loading control.

To further investigate the effect of NTRC on the reduction state of PQ pool in darkness, we measured the kinetics of Chl a fluorescence OJIP transients in dark-adapted leaves and in leaves pre-illuminated with far red light (FR) to fully oxidize the PQ pool. The difference in F/Fm at the J phase of the transient (F_J_, 3 ms after onset of illumination) between dark-adapted and pre-illuminated leaves is an indicator for the redox state of the PQ pool in darkness (Toth et al., 2007; Stirbet et al., 2014). As demonstrated in Fig. 4 and Suppl. Table S1, the OE-NTRC line had a significantly larger proportion of reduced PQ in darkness when compared to WT. The *ndho* mutant had a more oxidized PQ pool in darkness than WT, while there was no significant difference between *pgr5* and WT (Fig. 4G), suggesting that the NDH complex is the main CEF pathway contributing to dark-reduction of the PQ pool. Attribution of the elevated dark-reduction of PQ in OE-NTRC to enhanced activity of the NDH complex was further supported by the observation that the redox state of the PQ pool in darkness was more oxidized in the OE-NTRC *ndho* line in comparison to WT and OE-NTRC (Fig. 4G). OJIP transients also showed higher dark-reduction of PQ in *ntrc* mutant when compared to WT (Fig. 4), but it must be noted that the overall kinetics of the OJIP transient in *ntrc* differed considerably from the other lines, which may affect the interpretability of results from the pleiotropic *ntrc* knockout line.

### NTRC enhances the generation of proton motive force during dark-to-light transitions

Reduction of plastoquinone by the thylakoid NDH complex is known to be coupled to translocation of protons to the lumen, which contributes to the formation of *pmf* and enhances ATP synthesis (Strand et al., 2017a). We therefore investigated whether the generation of *pmf* during dark-to-light transitions and in plants illuminated with different light conditions is affected by deficiency or overexpression of NTRC. The *pmf*, conductivity of the thylakoid membrane to protons (*g*_H+_) and proportions of the *pmf* components ΔpH and ΔΨ were determined by measuring a difference in absorbance at 550 and 515 nm, also known as the electrochromic shift (ECS) (Cruz et al., 2005).

In WT, *pmf* was transiently elevated upon onset of illumination at growth light intensity, peaking after 15–20 seconds (Fig. 4) and coinciding with a decrease in *g*_H+_ (Fig. 5). The initial decrease in *g*_H+_ occurred despite rapid reduction of the gamma subunit of the ATP synthase (CF_1_*γ*), as after 20 s under growth light CF_1_*γ* was already fully reduced (Fig. 5) (Kramer et al., 1990). Under growth light intensity, another slight rise in *pmf* was observed after ca. 30–60 seconds in light, coinciding with P700 oxidation (Fig. 5 and Fig. 6). Subsequently *pmf* slowly decreased to a steady state value.

**Figure 5.**
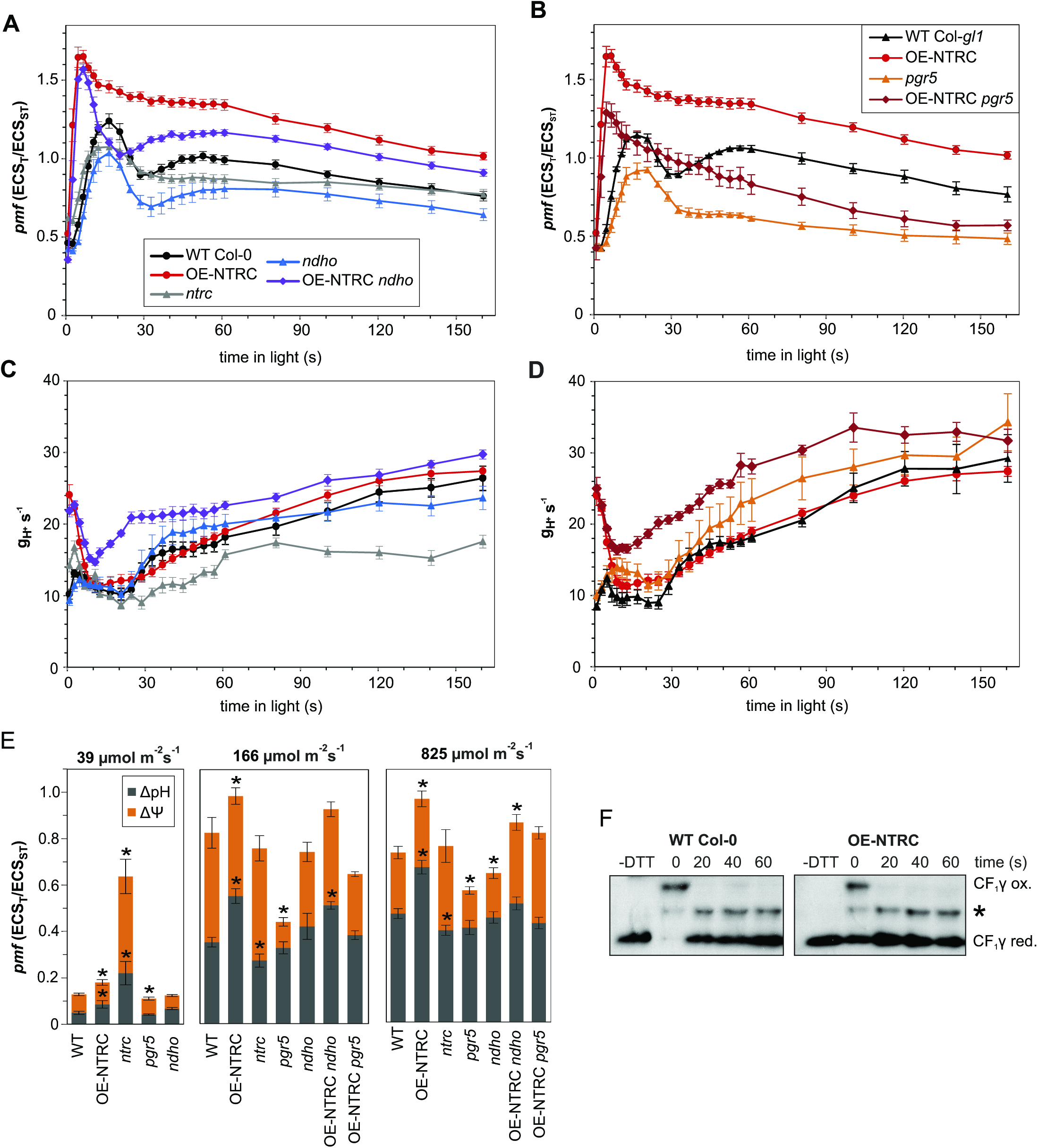
Generation of the proton gradient during dark-to-light transitions. **(A–B)** Proton motive force (*pmf*) at specific time points during transitions from dark to 166 μmol photons m^−2^s^−1^ actinic light (AL) in dark-adapted leaves of WT Col-0, OE-NTRC, *ntrc, ndho* and OENTRC *ndho* (A) and in WT Col-*gl1*, OE-NTRC, *pgr5* and OE-NTRC *pgr5* (B). The *pmf* was measured as light-induced change in the ECS signal (ECS_T_) and normalized with the magnitude of ECS induced by a 20 μs saturating single turnover flash administered prior to the onset of AL (ECS_ST_). Values are averages of measurements from 4 to 16 individual leaves ±SE. **(C–D)** Conductivity of the thylakoid membrane to protons (*g*_H+_), calculated as the inverse of the time constant of a first order fit to the decay of ECS during 250 ms dark intervals. **(E)** Total *pmf* and its partitioning to ΔpH and ΔΨ after 3 min illumination with low, growth or high light. **(F)** Mobility shift assays with MAL-PEG labelled protein extracts to determine the *in vivo* redox state of CF_1_*γ* subunit of the ATP synthase during first 60 seconds of dark-to-growth light transitions. * marks an unspecific band of unknown origin.

**Figure 6.**
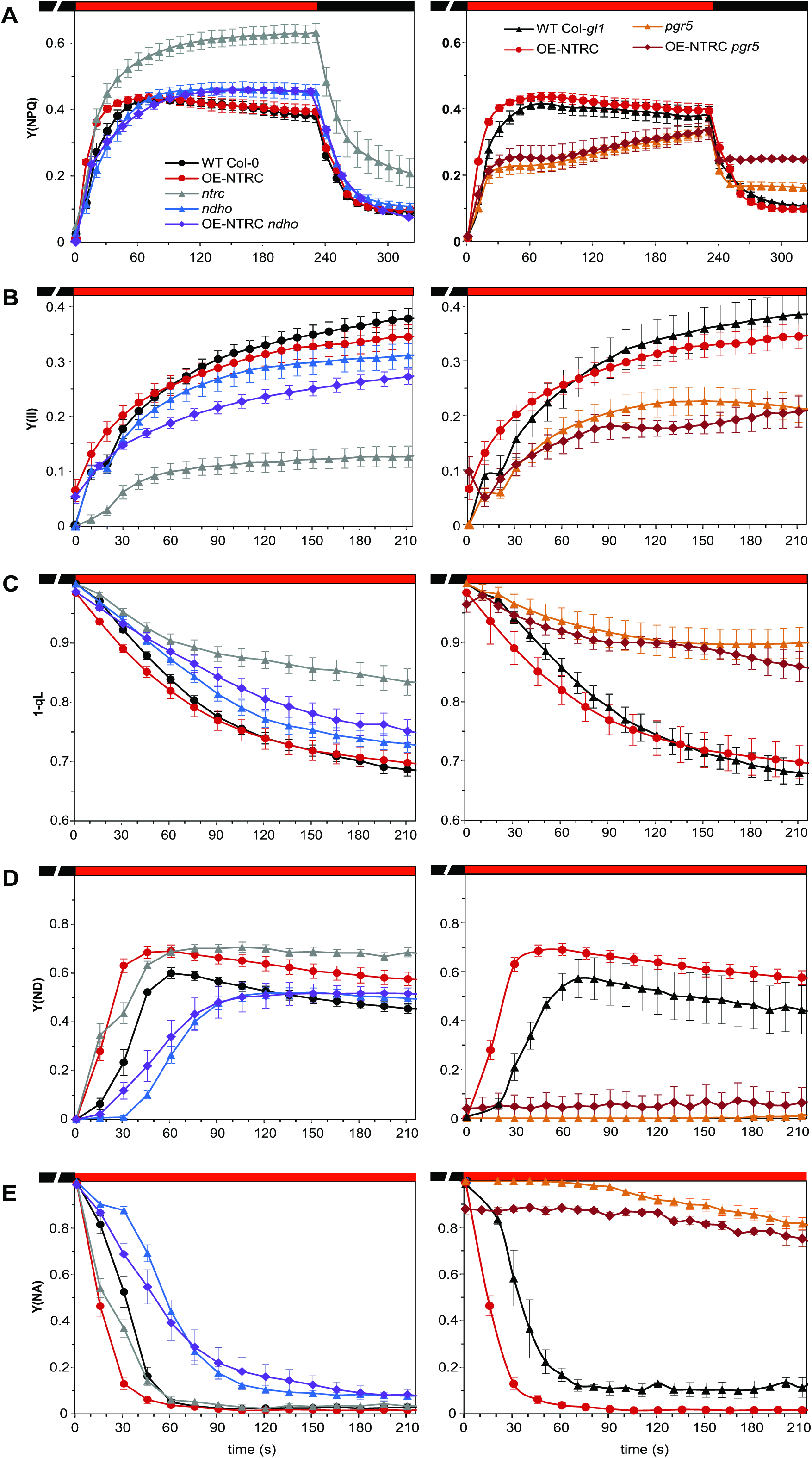
Photosynthetic parameters during dark-to-light transitions. **(A–E)** Induction of non-photochemical quenching (A), photosystem II quantum yield (B), redox state of the PQ pool (1-qL) (C), P700 oxidation (Y(ND)) (D) and PSI acceptor side limitation (Y(NA)) (E) were calculated from Chl a fluorescence and absorbance difference at 875 and 830 nm during transitions from dark to 166 μmol photons m^−2^s^−1^ actinic light in dark-adapted WT Col-0, *ntrc*, OENTRC, *ndho*, OE-NTRC *ndho*, WT Col-*gl1, pgr5* and OE-NTRC *pgr5* leaves. The graphs are averages of measurements from 4 to 9 individual leaves ±SE.

In OE-NTRC the initial *pmf* increase occurred already in a few seconds after the onset of illumination, reached a higher level and decreased more slowly than in WT (Fig. 5). While *g*_H+_ was drastically elevated in dark-adapted OE-NTRC leaves, it rapidly decreased to a level comparable to WT (Fig. 5). Both total *pmf* and the contribution of ΔpH to it were higher in OE-NTRC in all light intensities when compared to WT (Fig. 5). There was no significant difference between OE-NTRC and WT in *g*_H+_ under growth light illumination apart from the enhanced conductivity in dark-adapted leaves (Fig. 5) as shown previously (Nikkanen et al., 2016). Moreover, the PSII quantum yield (Fig. 6) increased only slightly during early photosynthetic induction and slightly but insignificantly decreased at steady state illumination in comparison to WT, while P700 oxidation was significantly enhanced (Fig. 6). These results strongly suggested that the increase in *pmf* in OE-NTRC derives from CEF. Despite a higher ΔpH under growth light intensity in OE-NTRC, we observed elevated NPQ only during the first minute after the dark-to-light transition (Fig. 6A). At steady state OE-NTRC had similar NPQ to WT.

In *ndho*, generation of *pmf* followed similar kinetics as in WT, but its level remained lower (Fig. 5). In comparison to WT, thylakoid proton conductivity was increased during the 30–60 s time period (Fig. 5) and NPQ induction was slightly delayed (Fig. 6A). In OE-NTRC *ndho*, however, an elevation of *pmf* occurred during transitions from dark to growth light similarly to OE-NTRC, except for a time period between 15–40 s after onset of illumination, where *pmf* was lowered in OE-NTRC *ndho* in comparison to OE-NTRC (Fig. 5). Upon dark-to light transitions the high initial thylakoid proton conductivity in OE-NTRC *ndho* decreased like in OE-NTRC, but after 10 s of illumination *g*_H+_ again rose more rapidly than in OE-NTRC (Fig. 5). As in *ndho*, NPQ induction was slightly delayed in OE-NTRC *ndho* (Fig. 6A). In the absence of the NDH complex, NTRC overexpression was also unable to enhance P700 oxidation during dark-to-growth light transitions, and OE-NTRC *ndho* suffered from increased PSI acceptor side limitation in comparison to WT (Fig. 6). This data indicates that the enhanced capacity of the stroma in OE-NTRC to pull electrons from PSI during dark-to-light transitions is dependent on the NDH-complex.

Increased activation of the NDH complex is not, however, sufficient to fully explain the elevated *pmf* in OE-NTRC, especially immediately after dark-to-light transitions and at steady state. Therefore, we also generated an NTRC overexpression line in the *pgr5* mutant background (OE-NTRC *pgr5*) whose NTRC expression level was comparable to OE-NTRC (Suppl. Fig. S1). In the *pgr5* mutant, *pmf* generation and NPQ induction at the onset of illumination were impaired in comparison to WT (Fig. 5 and Fig. 6). The secondary *pmf* rise was missing and the steady state level of *pmf* was lower than in WT, in part due to increased *g*_H+_ (Fig. 5). Absence of PGR5 did not alter the kinetics of *pmf* induction in plants overexpressing NTRC, but the magnitude of *pmf* remained lower (Fig. 5). At steady state, *pmf* in the OE-NTRC *pgr5* plants did not differ from *pgr5* plants (Fig. 5), while proton conductivity of the thylakoid membrane was drastically higher even in comparison to *pgr5* (Fig. 5). Moreover, PSI acceptor side limitation was only slightly alleviated in OE-NTRC *pgr5* in comparison to *pgr5* (6E). Interestingly, a high initial *g*_H+_ value and rapid generation of a *pmf* peak after onset of illumination were observed in OE-NTRC, OE-NTRC *ndho* and OE-NTRC *pgr5* plants but missing in WT, *ndho* and p*gr5* plants (Fig. 5). This demonstrates that the first rapidly-induced *pmf* peak in plants overexpressing NTRC is not caused by elevated CEF.

In *ntrc*, the initial *pmf* increase occurred with similar kinetics to WT upon onset of illumination at growth light intensity, but had a lesser magnitude (Fig. 5). The secondary *pmf* increase was absent in *ntrc* leaves. At steady state the *pmf* was comparable to WT, but contribution of ΔpH to total *pmf* was slightly diminished (Fig. 5). NPQ was however elevated (Fig. 6), implying active ΔpH-independent upregulation of NPQ. Decreased thylakoid conductivity to protons was observed in growth light intensity in *ntrc* (Fig. 5). Reduction of CF1*γ* is impaired in *ntrc* under low light but not in growth light (Nikkanen et al., 2016), implying that thylakoid proton conductivity is inhibited by other means. Not only in *ntrc*,but also in *pgr5* and OE-NTRC *pgr5* Q_A_ was maintained in a highly reduced state during dark-to-growth light transitions (Fig. 6). In *pgr5* and OE-NTRC *pgr5* this was due to impaired induction of NPQ (Fig. 6), lack of photosynthetic control at Cyt *b6f* and the consequential inability to oxidize PSI (Suorsa et al., 2012) (Fig. 6). In *ntrc*, however, high donor side limitation of PSI was measured in growth light despite high excitation pressure in the PQ pool (Fig. 6) and lowered PsaB content (Fig. 3), suggesting inhibition of electron transfer between the PQ pool and PSI, possibly at Cyt *b6f*. Lower and higher content of PGR5 and PTOX, respectively, (Fig. 3) may assist to relax excitation pressure in the PQ pool of *ntrc* plants.

### In fluctuating light NTRC overexpression enhances PSI yield in low light and represses thylakoid conductivity to protons upon transitions from low to high irradiance

Due to the recent suggestions implicating a particularly important role for NTRC under low and fluctuating light conditions (Nikkanen et al., 2016; Carrillo et al., 2016; Thormählen et al., 2017), we proceeded to investigate the *pmf* generation and photosynthetic electron transfer in OE-NTRC under these conditions. During transitions from dark to low light intensity, increased *pmf* formation was again observed in OE-NTRC, but the difference to WT was less dramatic than in growth light (Fig. 7). PSII yield was enhanced in OE-NTRC during the transition from dark to low light (Fig. 8), contributing to the *pmf* increase. P700 oxidation was also enhanced during dark to low light transitions (Fig. 9), while NPQ was decreased, despite higher ΔpH (Fig. 8). A high PSI yield was maintained throughout the low light periods in OE-NTRC due to low acceptor side limitation (Fig. 9). Notably, the overexpression of NTRC in *ndho* background reverted these changes observed in OE-NTRC plants to levels comparable to *ndho* knockout plants, except for the increased PSII yield during dark-to-low light transitions (Fig. 7, Fig. 8, Fig. 9), suggesting that enhanced activation of NDH-mediated CEF contributes to photosynthetic performance of OE-NTRC during dark to light transition and under low light.

**Figure 7.**
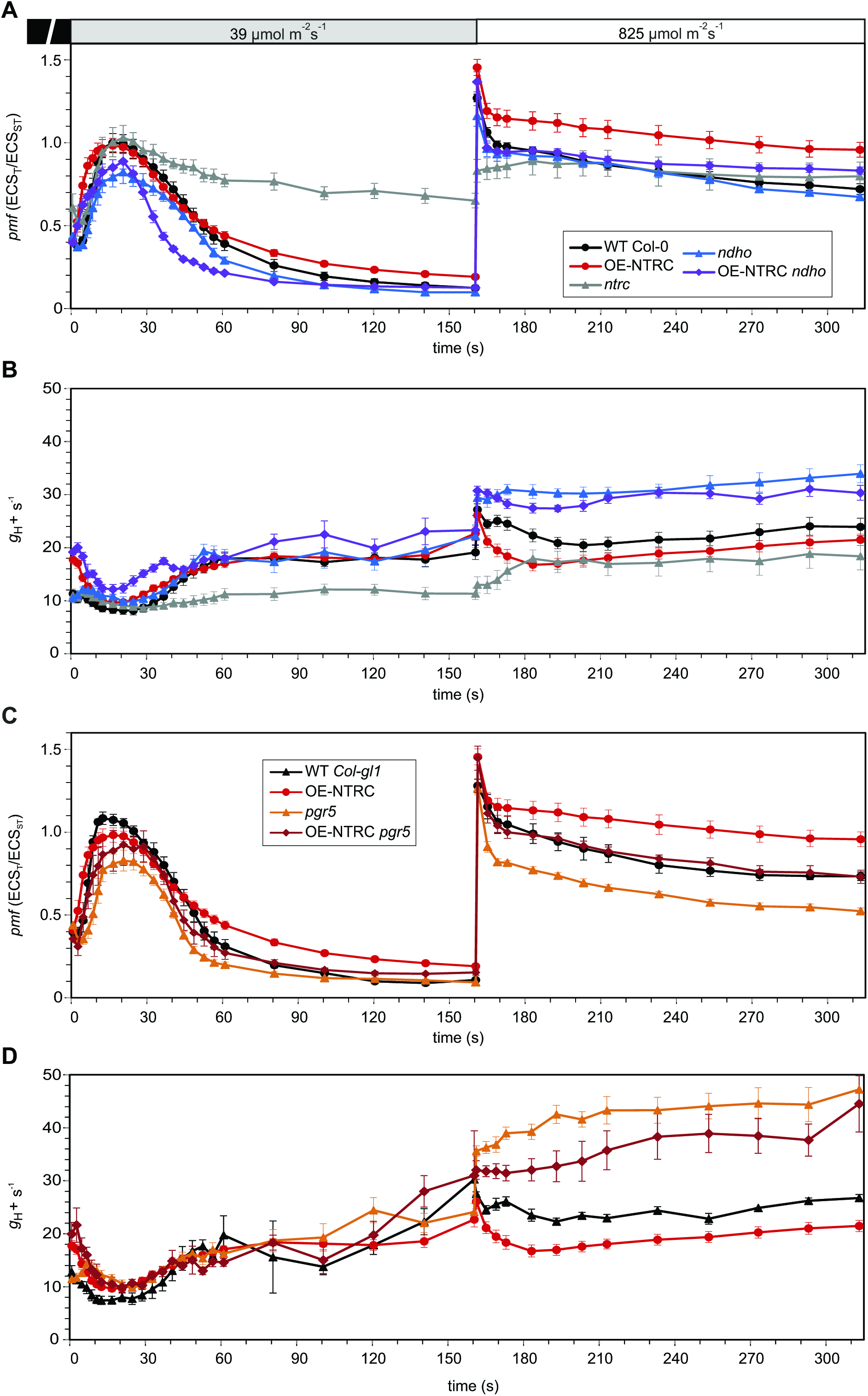
Formation and regulation the proton motive force during changes in light conditions. **(A–B)** The *pmf* (A) and proton conductivity of the thylakoid membrane(*g*_H+_) (B) at specific time points during transitions from darkness to low actinic light (39 μmol photons m^−2^s^−1^) and from low to high light (825 μmol photons m^−2^s^−1^) in dark-adapted leaves of WT Col-0, OE-NTRC, *ntrc, ndho* and OENTRC *ndho*. The *pmf* and *g*_H+_ were measured and calculated as explained in the legend for Figure 5. The graphs shown are averages of measurements from 4 to13 individual leaves ±SE. **(C–D)** The *pmf* and *g*_H+_ in dark-adapted leaves of WT Col-*gl1*, OE-NTRC, *pgr5* and OE-NTRC *pgr5*. The experiment was performed as explained in the figure legend for A–B. The graphs shown are averages of measurements from 3 to 13 individual leaves ±SE.

**Figure 8.**
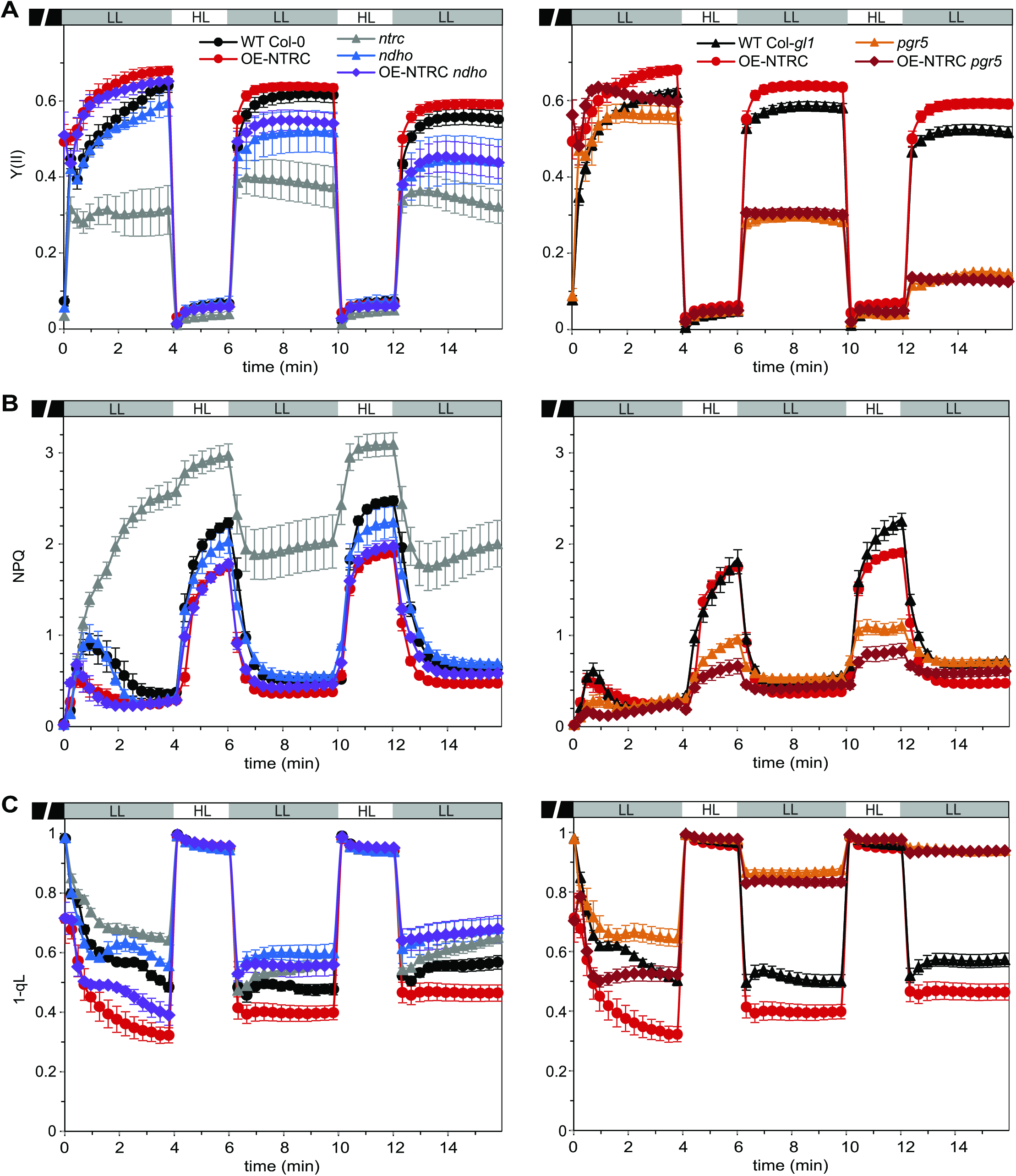
Analysis of Chlorophyll a fluorescence in fluctuating light. **(A-C)** PSII yield (Y(II)) (A), non-photochemical quenching (NPQ) (B) and redox state of the PQ pool (1-qL) (C) in light conditions fluctuating between periods of low actinic light (LL, 39 μmol photons m^−2^s^−1^) and high light (HL, 825 μmol photons m^−2^s^−1^) in WT Col-0, OE-NTRC, *ntrc, ndho*, OE-NTRC *ndho*, WT Col-*gl1, pgr5* and OE-NTRC *pgr5*. Five weeks old plants were dark-adapted for 30 min before measuring fluorescence from detached leaves. All values are averages of measurements from 3 to 10 individual leaves ±SE.

**Figure 9.**
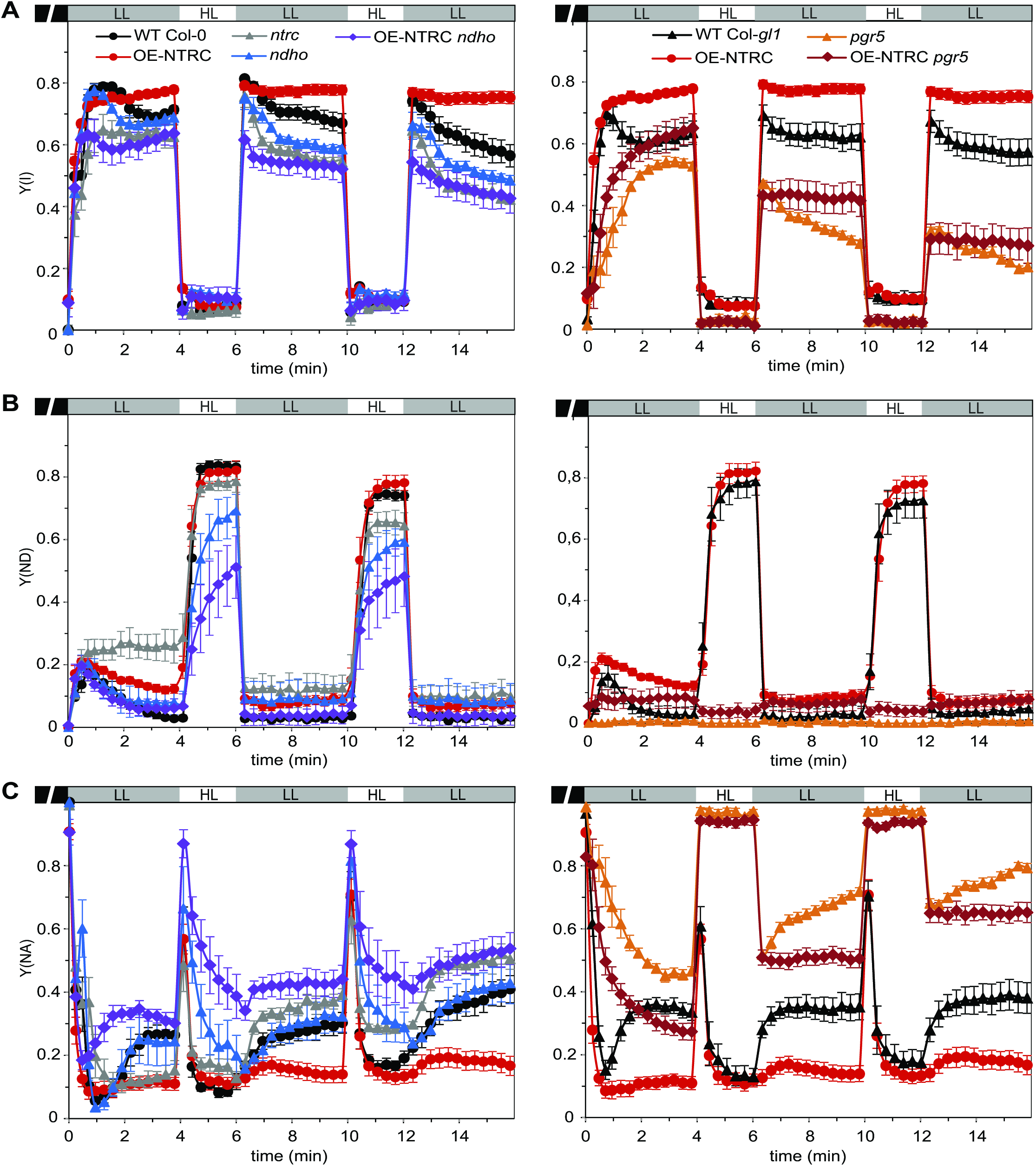
Analysis of P700 oxidation in fluctuating light. **(A-C)** PSI yield (Y(I)) (A), P700 oxidation (Y(ND)) (B) and PSI acceptor side limitation (Y(NA)) (C) in light conditions fluctuating between periods of low actinic light (LL, 39 μmol photons m^−2^s^−1^) and high light (HL, 825 μmol photons m^−2^s^−1^) in WT Col-0, OE-NTRC, *ntrc, ndho*, OE-NTRC *ndho*, WT Col-*gl1, pgr5* and OE-NTRC *pgr5*. Five weeks old plants were dark-adapted for 30 min before measuring fluorescence from detached leaves. All values are averages of measurements from 3 to 10 individual leaves ±SE.

During transitions from dark to low light, the *pgr5* and *ndho* mutants both showed impaired *pmf* generation during transitions from dark to low light, with most severe impairment occurring in *pgr5* at 30–60 seconds after the onset of low illumination. (Fig. 7). It is worth noting that this is the same time frame where transient NPQ and P700 oxidation are induced in WT but not in *pgr5* (Fig. 8 and Fig. 9). In contrast, *pmf* generation in OE-NTRC *pgr5* during dark-to-low and low-to-high light transitions was recovered to WT levels (Fig. 7) despite elevated *g*_H+_ especially in high light (Fig. 7), most likely due to enhanced activity of NDH-CEF. Slightly improved tolerance to light fluctuation was also observed in OE-NTRC *pgr5* in comparison to *pgr5* as the PSI yield was better maintained in low light following the high light periods (Fig, 9). Importantly, overexpression of NTRC improved the ability to oxidize PSI in low light even in the *pgr5* background (Fig. 9).

In all lines a transient *pmf* spike was observed upon the switch from low to high light intensity (Fig. 7). In WT, the proton conductivity of the thylakoid membrane decreased gradually (Fig. 7 and Fig. 8), but the decrease in *g*_H_+ was not due to oxidation of CF_1_*γ*, as it remained fully reduced in high light conditions (Suppl. Fig. S3). The decrease of *g*_H+_ was even stronger in OE-NTRC upon the shift from low to high light (Fig. 7). Despite elevated *pmf* (Fig. 7), NPQ induction was attenuated (Fig. 8). Overexpression of NTRC in *ndho* background decreased *pmf* generation during transitions from low to high light in comparison to the OE-NTRC line (Fig. 7), suggesting that enhanced activation of NDH-mediated CEF contributes to the high *pmf* in OE-NTRC in these conditions. OE-NTRC *ndho* showed increased steady-state *g*_H+_ under high irradiance similarly to *ndho* (Fig. 7).

The *pgr5* mutant was unable to oxidize P700 in high light (Fig. 9) or to decrease proton efflux from the lumen (Fig. 7), resulting in a loss of *pmf* (Fig. 7). The strong decrease of *g*_H+_ observed in OE-NTRC in high light (Fig. 7) disappeared in OE-NTRC *pgr5*, which lacks PGR5 (Fig. 7), suggesting that PGR5 contributes to the downregulation of thylakoid proton conductivity under these conditions. Recovery of *pmf* in high light through NTRC overexpression (Fig. 7) was not sufficient to induce NPQ (Fig. 8) or to control excess electron flow to PSI in *pgr5* background (Fig. 9). This supports the hypothesis that the PGR5 protein is directly required to support linear electron flow by inducing photosynthetic control at Cyt *b_6_f* (Suorsa et al., 2013; Tikkanen et al., 2015; Takagi and Miyake, 2018).

In *ntrc*, high steady state *pmf* under low light intensity (Fig. 7) was likely caused by impaired activation of the chloroplast ATP synthase and the Calvin-Benson cycle as previously reported (Nikkanen et al. 2016, Carrillo et al. 2016). Exceptionally high NPQ was recorded in the *ntrc* line, especially at low light (Fig. 8).

Concluding from Figures 5–9, it is evident that both the knockout and overexpression of NTRC had a distinct influence on the formation and regulation of *pmf* as well as on the induction of NPQ during transitions from dark to light and from low to high light through regulation of CEF activity and ATP synthase conductivity.

### Identification of CEF-related NTRC target proteins

Distinct effects of NTRC overexpression or deficiency on the post-illumination fluorescence rise (Fig. 2), the dark-reduction level of the PQ pool (Fig. 4) and generation of *pmf* as well as on the regulation of thylakoid conductivity during dark/light transitions and low/high light transitions (Fig. 5 and Fig. 7) suggested that NTRC may either directly or indirectly regulate CEF. In order to screen for potential targets of direct NTRC-mediated regulation, we performed co-immunoprecipitation (Co-IP) assays with an antibody against NTRC, and analyzed eluates from WT, *ntrc* and OE-NTRC total leaf extracts by mass spectrometry (MS). A full list of identified peptides is provided in Supplemental Dataset 1, while Suppl. Table S2 lists the 100 chloroplast proteins in order of abundance in WT and OE-NTRC eluates but absent in *ntrc* eluates. Among these proteins were several TRX interactors identified in previous studies (Balmer et al., 2003; Marchand et al., 2006; Hall et al., 2010), as well as established NTRC target proteins such as 2-Cysteine peroxiredoxins (Pérez-Ruiz et al., 2006) and enzymes involved in chlorophyll biosynthesis (Richter et al., 2013). Relevantly in the current context, several proteins involved in CEF around PSI were identified by the Co-IP/MS screening (Table 1). Most notably, five subunits of the thylakoid NDH complex; NdhH, Ndh48, NdhS, NdhU and NdhO, (in order of abundance in the Co-IP eluates) as well as PGR5 were identified as potential NTRC interactors. Intriguingly, all of the NDH subunits identified are located in close proximity to the proposed ferredoxin binding and oxidation site on the stromal side of the NDH complex (Yamamoto et al., 2011; Yamamoto and Shikanai, 2013; Peltier et al., 2016; Shikanai, 2016).

**Table 1.** Screening of putative NTRC target proteins in CEF pathways by Co-IP/MS. Only proteins of which at least two unique peptides were detected in WT / OE-NTRC eluates (+), and which were absent from *ntrc* eluates are included in the table. MW (kDa) indicates molecular weight of the protein, and #Cys the number of conserved cysteine residues (see Suppl. Table S3, Suppl. Table S4, and Suppl. Table S5). For a description of experimental procedures see Materials and methods, for a list of 100 chloroplast proteins detected in WT and/or OE-NTRC but not in *ntrc* eluates, see Suppl. Table 2, and for a full list of detected peptides see Suppl. Dataset 1.

| AGI code | Description | *ntrc* | WT | OE-NTRC | MW (kDa) | #Cys |
| --- | --- | --- | --- | --- | --- | --- |
| ATCG01110.1 | NdhH | - | + | + | 45.5 | 3 |
| AT1G15980.1 | Ndh48 (NDF1) | - | + | + | 51.0 | 3 |
| AT4G23890.1 | NdhS (CRR31) | - | + | + | 27.7 | 2 |
| AT1G74880.1 | NdhO | - | + | + | 17.6 | 0 |
| AT5G21430.2 | NdhU | - | + | - | 24.3 | 0 |
| AT2G05620.1 | PGR5 | - | + | - | 14.3 | 1 |

The potential interactions of NTRC with NdhS and PGR5 were further supported by positive results in bimolecular fluorescence complementation tests (BiFC). Co-expression of NTRC with both NdhS and PGR5 in *Nicotiana benthamiana* leaves resulted in YFP fluorescence that was strictly co-localized with chlorophyll autofluorescence, suggesting that it originated from the thylakoid membrane (Fig. 10). TRX-*m*1 interacted with PGRL1, while no interaction capability was detected between PGRL1 and NTRC (Fig. 10). Neither NdhS nor PGR5 interacted with TRX-*x* in BiFC, which was tested as a control (Suppl. Fig. S4).

**Figure 10.**
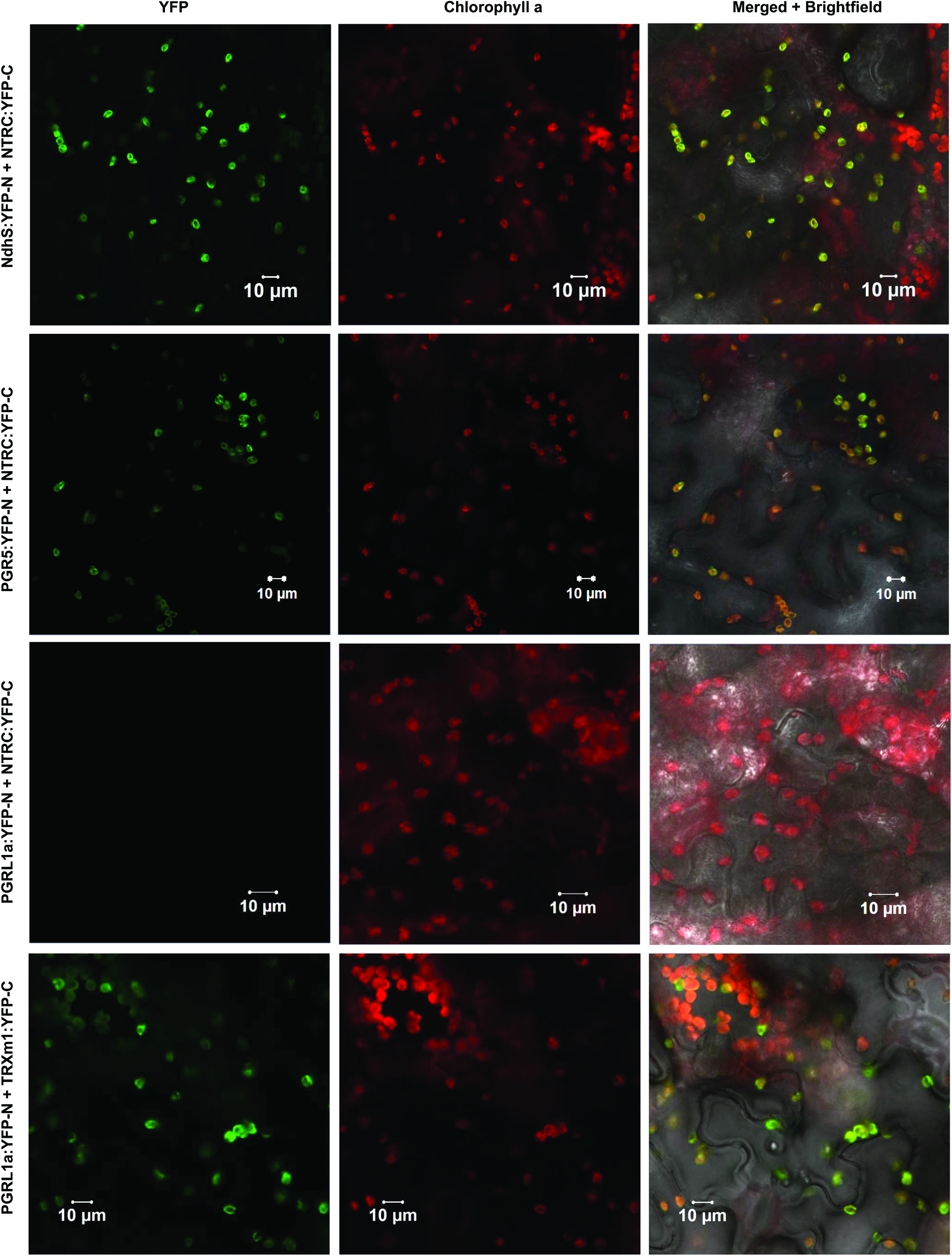
Bimolecular fluorescence complementation (BiFC) tests of *in planta* interactions between chloroplast TRXs and potential target proteins in cyclic electron flow. The left panel shows yellow fluorescent protein (YFP) fluorescence in green, the middle panel Chlorophyll a autofluorescence in red and the right panel a merged image of YFP, chlorophyll and brightfield images. YFP-N and YFP-C indicate expression of a fusion proteins including the N-terminal and C-terminal parts of YFP, respectively, in tobacco (*Nicotiana benthamiana*) leaves.

To assess the potential of the CEF-related proteins identified by Co-IP/MS (Table 1) and BiFC (Figure 10) to be targets of redox-regulation by TRXs, their amino acid sequences were analyzed for conserved cysteine residues. Of the NDH subunits identified as putative NTRC targets by Co-IP/MS, NdhS, NdhH, and Ndh48 contain cysteine residues that are conserved in angiosperms (Table 1, Suppl. Table S3, Suppl. Table S4, Suppl. Table S5), and therefore could in theory be subject to thiol regulation. NdhO and NdhU do not contain conserved cysteine residues, but they likely co-precipitate with the NTRC antibody because of their interactions with NdhH and NdhS subunits of the NDH complex, respectively (Shikanai, 2016).

PGR5 has been shown to form redox-dependent heterodimers with PGRL1 which have been proposed to be required for acceptance of electrons from ferredoxin and for reduction of PGRL1 (Hertle et al., 2013; Leister and Shikanai, 2013). The mature PGR5 polypeptide contains a single highly conserved cysteine residue (Munekage et al., 2002), which could hypothetically form an intermolecular disulfide with PGRL1 or some other partner, or be a target for S-nitrosylation or glutathionylation (Couturier et al., 2013; Zaffagnini et al., 2016). As direct determination of PGR5 redox state with the alkylation method was not feasible, we investigated if the redox state of PGRL1 is affected by NTRC deficiency or overexpression. PGRL1 contains six conserved thiol-sensitive cysteine residues that form inter- and intramolecular disulfides (Petroutsos et al., 2009; Hertle et al., 2013). We observed that PGRL1 was mostly oxidized in dark-adapted leaves, but underwent a transient reduction during approximately 60 seconds of illumination with growth light (Suppl. Fig. S3). This corresponds with the timescale of NPQ induction (Fig. 6), as well as with the transient increase in *pmf* and decrease in *g*_H+_ during dark-to-light transitions (Fig. 5). No significant difference to WT in PGRL1 reduction or protein content was detected in OE-NTRC (Suppl. Fig. S3, Fig. 3). PGR5 content of thylakoid membranes was, however, decreased in *ntrc* by 40% in comparison to WT (Fig. 3), in line with observations in a previous study (Yoshida and Hisabori, 2016).

## DISCUSSION

The role of CEF around PSI in plant acclimation to fluctuating light conditions has attracted great attention during the past 10 years. Importance of CEF likely relies in its capacity to maintain redox balance in chloroplasts upon fluctuations in light intensity and during dark-to-light transitions (Yamori et al., 2016; Suorsa et al., 2016; Strand et al., 2016b). Plastidial thioredoxin systems, on the other hand, are crucial regulators of chloroplast metabolic reactions in the stroma. It has, however, remained unclear whether thioredoxin-related redox regulation is also involved in the attainment of CEF-mediated redox balance upon exposure of plants to changing light intensities. To this end, we applied here an *in-vivo* approach to investigate whether NTRC contributes directly to regulation of CEF pathways in chloroplasts.

### Relevance of *ntrc* mutant in elucidating thioredoxin-dependent regulation of photosynthesis

The photosynthetic parameters measured for the *ntrc* knockout plants were not always in line with the results obtained with NTRC overexpressing plants (Fig. 2 and Fig. 4). In short photoperiod the *ntrc* plants have a highly pleiotropic phenotype (Fig. 3, Supl. Fig. 2) (Kirchsteiger et al., 2009; Pulido et al., 2010; Thormählen et al., 2015; Naranjo et al., 2016 Nikkanen et al., 2016; Pérez-Ruiz et al., 2017) that complicates the interpretation of results from the *ntrc* line. The high, slow-rising PIFR and increased dark-reduction of PQ in short-day grown *ntrc* (Fig. 2 and Fig. 3) were likely caused by a high NPQ (Fig. 8 and Suppl. Fig. S2) and its relaxation in darkness, lowered PSI content (Fig. 3) (Thormählen et al., 2015), and impaired stromal redox metabolism (Nikkanen et al., 2016; Pérez-Ruiz et al., 2017). Furthermore, we observed increased accumulation of reduced Fd, the substrate of the NDH complex, in *ntrc* (Fig. 2), which may be due to lower activity of CBC enzymes and consequent lower consumption of NADPH in carbon fixation (Nikkanen et al., 2016). Accordingly, a high NADPH/NADP^+^ ratio has been shown to activate CEF (Breyton et al., 2006; Okegawa et al., 2008). A high NADPH/NADP+ ratio together with high accumulation of reduced Fd may also explain the activation of NDH-dependent CEF by H_2_O_2_ (Strand et al., 2015; Strand et al., 2017), since oxidative treatment decreases the activation of redox-regulated CBC enzymes, and consequently the consumption of NADPH in illuminated chloroplasts. Accordingly, the *ntrc* mutant has a higher accumulation of H_2_O_2_ than WT (Suppl. Fig. S2 and Pulido et al., 2010), a higher NADPH/NADP+ ratio (Thormähälen et al., 2015), and lower activation states of CBC enzymes (Nikkanen et al., 2016; Pérez-Ruiz et al., 2017), which likely all contribute to the altered PIFR observed here. The PIFR was significantly diminished in *ntrc* plants grown in a longer photoperiod (Suppl. Fig. S2), where NPQ is lower (Suppl. Fig. S2) and the phenotype of *ntrc* has been shown to be less severe in terms of growth, chlorophyll content and efficiency of photochemistry (Lepistö et al., 2009; Lepistö et al., 2013). In *ntrc* electrons also accumulate in the PQ pool upon increases in light intensity (Fig. 6 and Fig. 8), although PSI is limited on the donor side (Fig. 6 and Fig. 9), indicating that electron transfer is limited between the PQ pool and PSI, possibly at Cyt *b6f*. Higher content of PTOX (Fig. 3) may assist to relax excitation pressure in the PQ pool of *ntrc* plants.

The pleiotropy described above for *ntrc* complicates the general interpretation of the results and makes it difficult to assess the contribution of NTRC to the regulation of photosynthesis when using the *ntrc* line. To avoid these obstacles we have constructed the OE-NTRC line, whose phenotype and development are not considerably dissimilar to WT (Toivola et al., 2013, Nikkanen et al., 2016) OENTRC in different backgrounds (*ntrc, ndho* and *pgr5*) provides a more reliable platform to examine the direct effects of NTRC on specific plastidial processes.

### NTRC in regulation of CEF

In-depth analysis of NTRC overexpression lines in respect to thylakoid functional properties provided strong evidence that NTRC indeed is involved in the regulation of CEF in the thylakoid membrane. Compelling support for NTRC-induced activation of the NDH complex was obtained by analyzing thylakoid CEF-related functions in NTRC overexpression lines made on the backgrounds of *ndho* (OE NTRC *ndho*) and *pgr5* (OE NTRC *pgr5*), incapable of performing NDH- and PGR-dependent CEF, respectively. Distinct effect of NTRC overexpression on the post-illumination rise of chlorophyll a fluorescence, the redox state of the plastoquinone pool in darkness as well as on the generation of the *pmf* and oxidation of P700 upon dark-to-light transitions and sudden increases in light intensity demonstrated the activating effect of NTRC on NDH-dependent CEF (Fig. 2, Fig. 3, Fig. 5, Fig. 6, and Fig 7). Higher activity of NDH-dependent CEF in plants overexpressing NTRC was shown not to be due to increased accumulation of the NDH complex or its substrate, because no increase in the accumulation of either NDH subunits or reduced Fd was detected in OE-NTRC plants in comparison to WT (Fig. 2 and Fig. 3). Furthermore, evidence for a direct effect of NTRC on the activation of NDH was obtained by identification of NDH subunits in a close proximity of the ferredoxin binding site as potential NTRC interactors (Table 1, Fig. 10). Although several NDH subunits were detected by Co-IP/MS, most likely only one or few of these subunits are genuine NTRC targets. The others likely co-precipitate with the NTRC antibody due to reciprocal interactions of the NDH subunits on the stromal side of the thylakoid membrane (Shikanai, 2016; Peltier et al., 2016). Existence of a thiol-regulated component in the ferredoxin binding site would provide a mechanism for dynamic control of the ferredoxin:plastoquinone oxidoreductase activity of the complex in response to fluctuations in light conditions. Redox-regulation of the NDH complex would allow rapid adjustment of *pmf* and nonphotochemical quenching as well as a maintenance of redox balance between the electron transfer chain and the electron sink capacity of stromal acceptors, most importantly the CBC. In high light, less active NDH could prevent the reverse function of the complex (i.e. oxidization of PQ to reduce ferredoxin and transfer of H^+^ from lumen to stroma) in conditions of high *pmf* and a reduced PQ pool (Strand et al., 2017a).

Notably, the overexpression of NTRC also affects the function of PGR5. Nevertheless, as discussed below, the effect is not necessarily related to the putative role of PGR5 in CEF. Our results, more likely, support the hypothesis (Avenson et al., 2005; Tikkanen et al., 2015; Kanazawa et al., 2017; Armbruster et al., 2017) that PGR5 is involved in control of the proton conductivity of the thylakoid membrane that, consequently, affects the generation of *pmf*.

### Regulation of *pmf* and the thylakoid redox balance via NDH and NTRC during changes in light conditions

Overexpression of NTRC caused elevated *pmf* under all light conditions, while no significant changes were observed in PSII quantum yield or thylakoid proton conductivity in comparison to WT (Fig. 5, Fig. 6, Fig. 7 and Fig. 8). These results strongly suggest that the elevation of *pmf* derives from enhanced CEF. Increased P700 oxidation during dark-to-light transitions in OE-NTRC was fully reverted in OE-NTRC *ndho*, and a lack of NDH also delayed the ability to oxidize P700 during the high light phases in fluctuating light (Fig. 6, Fig. 9). It is therefore evident that the NDH complex regulates the trans-thylakoid *pmf* as well as the redox balance between the electron transfer chain and the stroma, and this regulation is under the control of the stromal TRX systems, with our results suggesting a specific role for the NTRC system.

While our results demonstrate enhancement of the NDH-dependent CEF by NTRC overexpression (Fig. 2), earlier studies have revealed an inhibition of NDH-dependent CEF upon TRX-*m*4 overexpression and, conversely, an enhancement in *trxm4* mutants (Courteille et al., 2013). Thus, it is conceivable that the two chloroplast TRX systems regulate CEF in an antagonistic way, although it remains to be elucidated how such regulation might be mechanistically accomplished. We propose that in low light and upon sudden changes in the light intensity, NTRC is crucial for activation of the NDH-dependent CEF, while the TRX-*m4*-dependent inhibition of NDH-CEF requires higher light intensity or longer duration of illumination. Moderate to high-light illumination is required to fully activate the FTR-dependent TRX system (reviewed in Geigenberger et al. 2017) that possibly contributes to downregulation of NDH. In OE-NTRC the NDH-dependent CEF is constitutively active in light, which contributes to elevated *pmf* in all light intensities. Upon transition from dark to low light, there is less difference between OE-NTRC and WT in terms of *pmf* formation (Fig. 7), because in those conditions the NTRC-mediated activation of NDH occurs similarly in WT and OE-NTRC.

The NDH complex translocates protons from the stroma to the lumen not only via Cyt *b6f*, but also itself functions as a proton pump with a 2 H^+^/ e^−^ stoichiometry (Strand et al., 2017a). NDH-mediated CEF therefore contributes relatively more to *pmf* generation and consequently to ATP synthesis and NPQ induction than the PGR-dependent pathway. It has been postulated that the NDH complex is unlikely responsible for CEF during the early induction phase of photosynthesis, due to a low concentration of the complex in thylakoids in relation to the total PSI content (Joliot and Joliot, 2002). However, the NDH complex forms functional CEF-supercomplexes with PSI in stroma thylakoids (Peng et al., 2008), and a single NDH complex can bind up to six PSI complexes (Yadav et al., 2017), indicating that even a relatively low NDH content may have a significant impact on *pmf* generation.

### Contribution of TRX systems to PGR-related CEF pathway

Increased activation of NDH-CEF is not alone sufficient to explain all observed changes of *pmf* in OENTRC plants. When compared to WT, *pmf* remained elevated in OE-NTRC *ndho* during the first seconds of photosynthetic induction and at steady state in growth and high light (Fig. 5 and Fig. 7). These results could be explained by activation of PGR-dependent CEF as well in plants overexpressing NTRC. Stromal thiol redox state has been previously suggested to control the PGR-dependent CEF by a component that has a midpoint redox potential of −310 mV (Hertle et al., 2013; Strand et al., 2016a). It has also been proposed that *m*-type TRXs, with redox potentials between −357 and −312 mV (Collin et al., 2003; Yoshida et al., 2015), reduce an intermolecular disulfide in PGRL1 homodimers, and subsequently, the released monomeric PGRL1 may function as the ferredoxin-plastoquinone reductase (Hertle et al., 2013). Here we confirm the previously reported transient reduction of PGRL1 during dark-to-light transitions (Hertle et al., 2013), but NTRC overexpression does not intervene in the reduction (Suppl. Fig. S3). Moreover, TRX-*m*1 but not NTRC interacts with PGRL1 in BiFC (Fig. 10). Our results thus support the hypothesis that TRX-*m* is a primary reductant of PGRL1. Crosstalk between NTRC and FTR-dependent systems (Toivola et al., 2013; Thormählen et al., 2015; Nikkanen et al., 2016), and the interaction of NTRC with TRX-*m*1 in BiFC (Nikkanen et al., 2016), further support the interpretation that the activation of PGR-dependent CEF is indirectly increased in NTRC-overexpressing plants through enhancement of TRX-*m* reduction. This would also be in line with the steady state *pmf* increase observed in OE-NTRC *ndho* in comparison to WT (Fig. 5 and Fig. 7).

Alternatively, NTRC overexpression may affect the function of PGR5 in a way that is independent of its involvement in CEF. Redox regulation of PGR5 may occur to control its association with the ATP synthase during dark-to-light and low-to-high light transitions, and thereby allows inhibition of the ATP synthase in an unknown mechanism, as suggested earlier (Avenson et al., 2005; Tikkanen et al., 2015; Kanazawa et al., 2017; Armbruster et al., 2017). Such a mechanism would result in acidification of the lumen and induction of NPQ, allowing dissipation of excess excitation energy from the electron transfer chain until CBC is activated. This hypothesis is supported by the impaired abilities of *pgr5* and OE-NTRC *pgr5* to control the activity of the ATP synthase at early stages of dark-light transitions and upon transitions to high light intensities (Avenson et al., 2005, Fig. 5 and Fig. 7). Furthermore, the elevated NTRC content in leaves caused decreased thylakoid proton conductivity upon increases in light intensity (Fig. 7), suggesting that NTRC controls the PGR5-dependent down-regulation of proton efflux from lumen. This is supported by the identification of PGR5 as a potential NTRC interactor (Table 1, Fig. 3, Fig. 10).

The initial strong *pmf* increase in OE-NTRC after onset of growth light illumination was evident also in both the OE-NTRC *ndho* and OE-NTRC *pgr5* plants (Fig. 5) indicating that this *pmf* peak is not caused by CEF. More likely, the initial *pmf* results from dark-activation of the CBC enzymes in plants overexpressing NTRC (Nikkanen et al. 2016), which provides an enhanced ability of the stroma to accept electrons from the PETC upon dark-to light transition and consequently enhances proton pumping to the lumen.

### Cooperative regulation of photosynthetic electron transfer and carbon fixation by chloroplast thioredoxin systems

Light-dependent reductive activation of the ATP synthase, the CBC and the NADP-malate dehydrogenase (NADP-MDH) by TRXs has been well established for several decades (reviewed in (Buchanan, 2016). More recently, knowledge of TRX-mediated control has been extended to various regulatory and photoprotective mechanisms of photosynthesis, including regulation of state transitions (Rintamäki et al., 2000; Shapiguzov et al., 2016), NPQ (Hall et al., 2010; Brooks et al., 2013; Naranjo et al., 2016; Da et al., 2017) and CEF (Courteille et al., 2013; Hertle et al., 2013; Strand et al., 2016a). We propose here a model, comprising a cooperative function of the two chloroplast TRX systems with distinct reductants and redox potentials that allows the maintenance of redox balance between the two photosystems and stromal metabolism during fluctuations in light conditions. This is achieved through dynamic regulation of the activities of the ATP synthase, NPQ, the NDH complex, PGRL1/PGR5 as well as the LHCII kinase STN7 by reversible thiol modifications. We propose a specific role for NTRC in regulating NDH-CEF, the ATP synthase and CBC enzymes in low light, dark-to-light transitions and during sudden increases in light intensity, as schematically depicted in Fig. 11.

**Figure 11.**
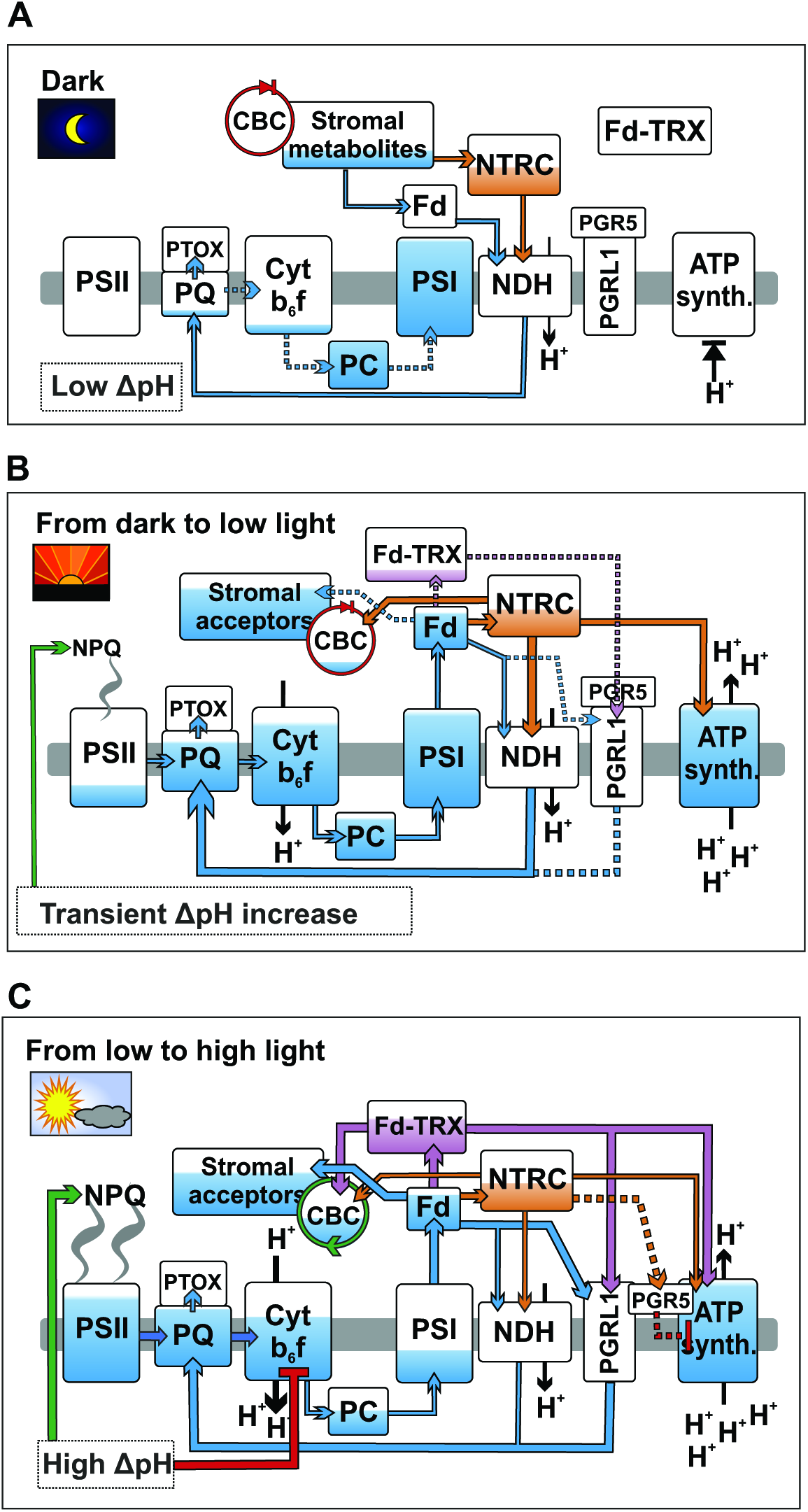
A schematic model of the role of chloroplast TRX systems in regulating CEF and the *pmf* during dark-to-light transitions and fluctuations in light intensity. **(A–C)** Dark-adapted leaves (A), transition from dark to low light (B) and transition from low to high light (C). Blue color indicates the approximate reduction levels of different photosynthetic redox components based on data in the current paper (Figures 1, 2, 8, and 9) as well as other reports (Yoshida and Hisabori, 2015; Nikkanen et al., 2016; Schreiber, 2017; Schreiber et al., 2017). Green and red arrows indicate activating and inhibitory effects, respectively, while orange color represents the thiol regulation by NTRC and purple by the Fd-TRX system. Thicker lines depict stronger effect than thin and dotted lines. For details see the text. The arrow to NPQ refers to induction of the qE component of NPQ due to acidification of the lumen (Demmig-Adams et al., 2012). The arrows representing reduction of PQ by the NDH complex and PGRL1 have been drawn through the lumen only to increase clarity of the illustration.

In darkness, a proportion of NDH complexes in the thylakoid membrane is activated by NTRC, and moderate chlororespiration from NDH to PTOX occurs. Due to an inactive ATP synthase and proton pumping activity of NDH, a weak proton gradient over the thylakoid membrane is established. Redox-regulated CBC enzymes are inactive, causing PC and P700 to be reduced (Schreiber, 2017) due to lack of electron acceptors in the stroma. In OE-NTRC, chlororespiration via NDH to the PQ pool is elevated due to increased amount of active NDH complexes. This leads to increased protonation of the lumen and higher reduction of the PQ pool. Proportions of the ATP synthase and CBC enzyme pools are activated due to high NTRC content.

Upon transition from dark to low light, the ATP synthase pool is fully and the CBC enzyme pool partially reduced by NTRC in WT plants. Delay in activation of the CBC enzymes causes, however, reduction of the PQ pool due to scarcity of stromal acceptors that limits electron transfer. NTRC contributes to activation of NDH-dependent CEF, which alleviates electron pressure at PSI and transiently increases *pmf* and induces NPQ. In OE-NTRC, P700 and PC are effectively oxidized upon onset of low illumination, as the acceptor side limitation is negligible due to fully active NDH-dependent CEF, ATP synthase and redox-activated CBC enzymes. This results in an elevated ΔpH and faster induction of NPQ in comparison to WT at the initial phase of illumination. At steady state NPQ is lower than in WT despite of high ΔpH, suggesting downregulation of NPQ by thioredoxin via a ΔpH-independent mechanism, as reported previously by Brooks et al. (2013).

When a leaf is shifted from low to high irradiance, both TRX systems become fully active, and the CBC enzymes as well as the PGR-dependent CEF are fully activated. NTRC affects PGR5-dependent inhibition of the ATP synthase, which contributes to accumulation of protons in the lumen. Consequently, NPQ and down-regulation of electron transfer at Cyt *b6f* are induced. Electrons are effectively pulled from PSI, and the donor side is limiting electron transfer. In OE-NTRC, increased reduction of PGR5 likely leads to stronger down-regulation of the ATP synthase. This, together with proton pumping by constantly active NDH and possibly through increased TRX-*m*-mediated activation of the PGR-dependent pathway, results in high *pmf*. NPQ is however lower than in WT due to ΔpH-independent downregulation of NPQ by overexpressed NTRC.

## CONCLUSIONS

In the present paper, we have shown that the chloroplast NADPH-dependent thioredoxin system (NTRC) stimulates cyclic electron flow around PSI by activating the thylakoid NDH complex. We propose that NTRC-mediated activation of CEF is particularly important during fluctuations in light intensity and in low light in order to keep redox balance between photosynthetic electron transfer chain in the thylakoid membranes and stromal metabolism. Interactions assays suggest that a TRX-target exists close to the ferredoxin binding and oxidation site of the NDH-complex, and possibly involves the NdhS subunit. We further suggest that NTRC regulates photosynthetic redox poise by promoting PGR5-dependent downregulation of the activity of the ATP synthase upon transitions from low to high light intensity. This results in acidification of the lumen, which is needed for induction of photoprotective mechanisms.

## MATERIALS AND METHODS

### Plant material and growth conditions

Experiments have been done with *Arabidopsis thaliana* wild type (WT) lines of the Columbia ecotype (Col-0 and Col-*gl1*), and with the following transgenic lines: NTRC overexpression line (Toivola et al., 2013), T-DNA knockout mutants of NTRC (At2g41680, SALK_096776) (Lepistö et al., 2009), *ndho* (At1g74880, SALK_ 068922) (Rumeau et al., 2005) and STN7 (AT1G68830, SALK_073254) (Bellafiore et al., 2005) as well as the *pgr5* mutant (AT2G05620) (Munekage et al., 2002). The plants were grown in a photoperiod of 8 h light / 16 h darkness at 23 °C under 200 μmol of photons m^−2^◻s^−1^ for all experiments except for the measurements shown in Suppl. Fig. S2, for which plants were grown in a 12 h /12 h photoperiod under 130 *μ*mol m^−2^◻s^−1^. Wild type tobacco (*Nicotiana benthamiana*) plants used in BiFC tests were grown under 130◻*μ*mol photons m^−2^◻s^−1^ at 23◻°C in a 16◻h light/8◻h dark photoperiod. The OE-NTRC *ndho* and OE-NTRC *pgr5* lines were generated by *Agrobacterium tumefaciens* and floral dipping –mediated transformation of the *ndho* knockout and *pgr5* mutant lines, respectively, with the NTRC overexpression construct as described previously (Toivola et al., 2013). The OE-NTRC *ndho* and OE-NTRC *pgr5* plants used in the experiments were heterozygous T2 generation plants that were selected on agar plates with 0.5X Murashige-Skoog medium (MS) (Murashige and Skoog, 1962) and 50 μg/ml kanamycin. The plants were subsequently transferred to soil and grown in a 8 h light / 16 h darkness photoperiod at 23 °C under 200 μmol of photons m^−2^◻s^−1^ for four weeks before usage in the experiments. As control, OE-NTRC plants were similarly selected on kanamycin-containing plates while WT Col-0 and WT Col-*gl1* (ecotype of the *pgr5* mutant) plants were grown on 0.5X MS-agar plates without antibiotics for an equivalent time.

### Determination of H_2_O_2_ content in leaves

The hydrogen peroxide content in leaves was estimated by staining with diaminobenzidine (DAB), as previously described in Lepistö et al. (2013). Detached leaves from 4-weeks-old WT, *ntrc* and OENTRC plants were incubated overnight in darkness in 0.1 mg ml^−1^ solution of diaminobenzidine (DAB; Sigma-Aldrich) (pH 3.8), after which the leaves were illuminated with either 40 or 200 μmol photons m^−2^◻s^−1^ for 1h. Chlorophyll was then bleached by incubating the leaves in ethanol and subsequently photographed. Image J software (Schneider et al., 2012) was used to quantify the intensity of the staining.

### Measurement of Chlorophyll a fluorescence and P700 and Fd redox changes

The post-illumination chlorophyll a fluorescence rise (PIFR) was measured from detached leaves with the Multicolor-PAM fluorometer (Walz). A 480 nm measuring beam at an intensity of 0.2 *μ*mol photons m^−2^◻s^−1^ was used to measure fluorescence changes after illumination of dark-adapted (30 min) leaves with 67 *μ*mol photons m^−2^◻s^−1^ of white actinic light for 500 seconds, with saturating pulses of 800 ms (10000 *μ*mol photons m^−2^◻s^−1^) in the beginning and at 400 s to determine Fm and Fm’. The actinic light was then switched off and the changes in chlorophyll a fluorescence in the dark were observed for 300 s. A 10 s pulse of far red light was then given to fully oxidize the PQ pool, and the subsequent re-reduction PQ pool was detected through a rise in Chl fluorescence.

The OJIP transients were recorded with the Multicolor-PAM from dark-adapted (30 min) leaves and from leaves pre-illuminated with far red light (intensity setting 15) for 6 s, according to the method described by (Toth et al., 2007). A saturating pulse of 3000 *μ*mol photons m^−2^◻s^−1^ and measuring light at 440 nm were used in the measurements.

The Dual-PAM-100 was used to simultaneously record the Chl a fluorescence and P700-dependent difference in absorbance at 875 and 830 nm during transitions from dark to 166 μmol photons m^−2^ s^−1^ (Fig. 6) and during a light regime, where a 620 nm AL fluctuates between 39 and 825 μmol photons m^−2^ s^−1^ (Fig. 8 and Fig. 9). Saturating pulses were administered at 10 or 15 s intervals for the measurements in Fig. 6 and at 15 s intervals for the first minute after onset of illumination, and at 20 s intervals thereafter for Fig. 8 and Fig. 9. Because *ntrc* leaves are very small in size and low in chlorophyll content, it was in some cases necessary to record from two or three leaves simultaneously to obtain a P700 signal of sufficient quality. The parameters shown were calculated with the Dual-PAM-100 software as described by Bilger and Björkman (1990), Klughammer and Schreiber (2008a, 2008b) and Kramer et al., (2004).

For determination of Fd redox state, the Dual/Klas-NIR (Walz) spectrometer was used to record the four absorbance differences between 785 and 840, 810 and 870, 870 and 970, as well as 795 and 970 nm, from which the redox changes of P700, PC and Fd were deconvoluted as described by Klughammer and Schreiber (2016) and Schreiber (2017). A similar illumination and post-illumination regime was used as described above for the measurement of PIFR, with the exception that dark-adapted leaves were illuminated with 61 μmol photons m^−2^ s^−1^ of 630 nm instead of white actinic light. Measured Fd redox changes were then normalized to the level maximal Fd reduction, which was determined according to Schreiber and Klughammer (2016).

### Measurement of electrochromic shift (ECS)

In order to measure the magnitude and kinetics of *pmf* formation, changes in the electrochromic shift (ECS, P515) signal were recorded with the Dual-PAM-100 and the P515/535 accessory module (Walz) (Schreiber and Klughammer, 2008, Klughammer et al., 2013). A dual beam difference signal between 550 and 515 nm was used to avoid distortion of results by scattering effects. A measuring light at a 2000 Hz pulse frequency was used in all ECS measurements. For the dark-to-light and low-to-high light transition measurements in Fig. 5 and Fig. 7, plants were first dark-adapted for 30 min. A single-turnover saturating flash (20 μs) of 14000 μmol photons m^−2^s^−1^ was then applied to obtain ECS_ST_, a maximum absorbance change value that was used to normalize all results to account for differences in leaf thickness and chlorophyll content between individual leaves and lines (Kramer and Crofts, 1989). The obtained values of ECS_ST_ were in good correlation with the differences in chlorophyll content in OE-NTRC and *ntrc* lines reported previously (Toivola et al., 2013). In order to distinguish the light-induced ECS change (ECS_T_) from signal drift and baseline change, dark intervals of 250 ms were applied at the following time points after the onset of AL illumination: 0.8; 2.7; 4.7; 6.7; 8.7; 10.7; 12.7; 16.7; 20.7; 24.7; 28.7; 32.7; 36.7; 40.6; 44.7; 48.7; 52.7; 56.7; 60.7; 80.6; 100.5; 120.5; 140.5 and 160.5 s after onset of illumination. Additionally, during the shift from low to high irradiance (Fig. 7), dark intervals were applied at 1.1; 5.1; 9.1; 13.1; 23.1; 33.1; 43.1; 53.1; 73.1; 93.1; 113.1; 133.1 and 153.1 s after the increase in light intensity. ECS_T_ was calculated as the difference between total ECS in light and an Y_0_ value obtained from the first-order exponential fit to the decay kinetics of the ECS signal during a dark interval. Total *pmf* was then calculated as ECS_T_/ECS_ST_. The g_H_+ parameter, describing thylakoid membrane conductivity to protons, was calculated as the inverse of the time constant of a first-order exponential fit to ECS decay kinetics during a dark interval (Cruz et al., 2001; Avenson et al., 2005; Cruz et al., 2005). Partitioning of total *pmf* to its components ΔpH and ΔΨ was determined from the light-off response of the ECS signal (Cruz et al., 2001) after 3 min illumination, also using the Dual PAM ECS module as described by Schreiber and Klughammer (2008). Same settings were used for determination of *pmf* partitioning as for the dark-to-light and low-to-high light transition measurements.

### Protein extraction, alkylation of thiols and SDS-PAGE

Proteins and thylakoids were isolated as previously described (Lepistö et al., 2009), while chlorophyll content was determined according to (Porra et al., 1989) and protein content with the Bio-Rad Protein Assay kit. For determination of the redox states of TRX-regulated proteins, leaf proteins were precipitated with trichloroacetic acid (TCA) and free thiols in proteins alkylated with N-ethylmaleimide (NEM, Sigma-Aldrich). After alkylation protein disulfides were reduced with dithiothreitol (DTT, Sigma-Aldrich) and subsequently produced thiols were alkylated with methoxypolyethylene glycol maleimide M_n_ 5000 (MAL-PEG, Sigma-Aldrich) as described earlier (Nikkanen et al., 2016). Sodium dodecyl sulfate polyacrylamide gel electrophoresis (SDS-PAGE) and immunoblotting was performed as reported in (Nikkanen et al., 2016). For running the MAL-PEG samples pre-cast 4-20% Mini-PROTEAN TGX gels (Bio-Rad) were used, except for the gel in Fig. 1B, Suppl. Fig. S1A and Suppl. Fig. S3A, where a 12% polyacrylamide gel was used. PVDF membranes were probed with antibodies raised against NTRC (Lepistö et al., 2009), D1 (Research Genetics, Inc (Thermo Fisher)), PsaB (Agrisera, AS10 695), Cyt *f* (kindly provided by L. Zhang), PTOX (kindly provided by M. Kuntz), NdhH (Agrisera), NdhS (Agrisera), CF_1_*γ* (Agrisera, AS08 312), PGRL1 (Agrisera, AS10 725), PGR5 (Agrisera) or phosphothreonine (P-Thr) (New England Biolabs). Membranes were then treated with a horseradish peroxidase (HRP)-conjugated goat anti-rabbit secondary antibody (Agrisera, AS09 602) for 2◻h. All immunoblots shown are representative of at least three biological replicates. Quantifications of protein content shown in Fig. 3 were performed using the ImageJ software (Schneider et al., 2012) and normalized according to the intensity of Li-Cor Revert Total Protein Stain. Statistical significance was determined using two-tailed Student’s T-tests for unequal variances with p-values below 0.05 interpreted as statistically significant.

### Co-immunoprecipitation and Mass spectrometry

For co-immunoprecipitation (Co-IP), WT, *ntrc* and OE-NTRC leaves were frozen in liquid N2, lysed in Pierce IP Lysis buffer containing 1 % NP-40 detergent (Thermo-Fisher), and immunoprecipitated in a resin containing NTRC-specific antibody using the Pierce Co-IP kit (Thermo-Fisher) with an affinity-purified NTRC-specific antibody, as described previously (Nikkanen et al., 2016). Co-IP eluates were denatured and purified by SDS-PAGE in a 6% acrylamide gel with 6 M urea, subjected to in-gel tryptic digestion and the extracted peptides analyzed with the Q Exactive Hybrid Quadruple-Orbitrap mass spectrometer (Thermo-Fisher Scientific) in DDA mode as previously described (Trotta et al., 2016). MS/MS spectra were analyzed with an in-house installation of Mascot (v.2.4) (Matrix Science) search engine and analyzed with Proteome Discoverer (v.1.4) Software (Thermo Scientific), restricting the search to the non-redundant database TAIR10 supplemented with most common laboratory contaminants (Trotta et al., 2016). Peptides were validated by Decoy Database Search, with target false discovery rates (FDR) set to be below 0.01 (strict) or below 0.05 (relaxed).

### BiFC tests

Bimolecular fluorescence complementation tests (BiFC) were performed as described in (Nikkanen et al., 2016). For the current study, coding sequences of PGR5, PGRL1a and NdhS obtained from Arabidopsis Biological Resource Center (ABRC) were cloned into pSPYNE-35S and pSPYCE-35S binary vectors (Walter et al., 2004), and the resulting constructs were checked by sequencing. Primer sequences used for cloning are listed in Suppl. Table S6. Imaging of YFP and chlorophyll autofluorescence from *N. benthamiana* leaves infiltrated with *Agrobacterium tumefaciens* strain GV3101 carrying the appropriate binary vectors was performed with a Zeiss LSM780 laser scanning confocal microscope at 3 days after infiltration. The negative result between PGRL1:YFP-N and NTRC:YFP-C also serves as a negative control.

### Multiple alignment of amino acid sequences

Amino acid sequences of NdhH, Ndh48, NdhS, NdhJ and Ndh45 in *Arabidopsis thaliana* and, as available, in *Populus trichocarpa, Vitis vinifera, Glycine max, Solanum lycopersicum, Oryza sativa, Sorghum bicolor, Brachypodium distachion, Physcomitrella patens, Selaginella moellendorffii* and *Synechocystis PCC 6803* were obtained from the UniProtKB database and aligned with the Clustal Omega 1.2.4 online alignment tool (Sievers et al., 2011) using default settings.

### Accession Numbers

The Arabidopsis Genome Initiative locus identifiers (AGI) used in this paper are listed in Table 1, Suppl. Table S2, Suppl. Table S3, Suppl. Table S4, Suppl. Table S5, Suppl. Table S6 and Suppl. Dataset 1.

## Supplemental Material

Supplemental Figure S1. NTRC redox state in different light conditions in OE-NTRC and level of NTRC expression in NTRC overexpression lines.

Supplemental Figure S2. Post-illumination fluorescence rise in dark-adapted WT, OE-NTRC and *ntrc* plants grown in a 12 h/12 h photoperiod, and accumulation of H_2_O_2_ in illuminated leaves.

Supplemental Figure S3. *In vivo* redox states of PGRL1 and CF1*γ* during changes in light conditions.

Supplemental Figure S4. Bimolecular fluorescence complementation (BiFC) tests between TRX-*x* and NdhS and PGR5.

Supplemental Table S1. Parameters determined from OJIP transients.

Supplemental Table S2. Screening of putative NTRC interactors by Co-IP/MS

Supplemental Table S3. Multiple alignment of NdhS amino acid sequences.

Supplemental Table S4. Multiple alignment of NdhH amino acid sequences.

Supplemental Table S5. Multiple alignment of Ndh48 amino acid sequences.

Supplemental Table S6. Primers used for cloning of BiFC constructs.

Supplemental Dataset 1. MS/MS identification of peptides in Co-IP eluates.

## Author contributions

L.N. and E.R. designed the research, L.N., J.T., and A.T. performed the research, L.N., J.T., A.T., M.T., and E.R. analyzed the data, L.N. and E.R. wrote the article with input from E-M.A., M.T., A.T., M.G.D., and J.T.

## ACKNOWLEDGEMENTS

We thank Jesse Ojala for assistance with the experiments, Esa Tyystjärvi for the advice for PIFR measurements, Sari Järvi and Marjaana Suorsa for advice on the use of antibodies, Alexandrina Stirbet for expert advice on measurement of OJIP transients, and Mika Keränen, Kurt Ståhle and Tapio Ronkainen for technical assistance.

## LITERATURE CITED

Allorent, G., Osorio, S., Vu, J.L., Falconet, D., Jouhet, J., Kuntz, M., Fernie, A.R., Lerbs-Mache, S., Macherel, D., Courtois, F., and Finazzi, G. (2015). Adjustments of embryonic photosynthetic activity modulate seed fitness in Arabidopsis thaliana. New Phytol. 205, 707–719.

Armbruster, U., Correa Galvis, V., Kunz, H.H., and Strand, D.D. (2017). The regulation of the chloroplast proton motive force plays a key role for photosynthesis in fluctuating light. Curr. Opin. Plant Biol. 37, 56–62.

Armbruster, U., Carrillo, L.R., Venema, K., Pavlovic, L., Schmidtmann, E., Kornfeld, A., Jahns, P., Berry, J.A., Kramer, D.M., and Jonikas, M.C. (2014). Ion antiport accelerates photosynthetic acclimation in fluctuating light environments. Nature Comm. 5, 5439.

Avenson, T., Cruz, J., Kanazawa, A., and Kramer, D. (2005). Regulating the proton budget of higher plant photosynthesis. Proc. Natl. Acad. Sci. U. S. A. 102, 9709–9713.

Bailey, S., Walters, R.G., Jansson, S., and Horton, P. (2001). Acclimation of Arabidopsis thaliana to the light environment: the existence of separate low light and high light responses. Planta 213, 794–801.

Balsera, M., Uberegui, E., Schürmann, P., and Buchanan, B.B. (2014). Evolutionary Development of Redox Regulation in Chloroplasts. Antioxid. Redox Signal. 21, 1327–1355.

Bellafiore, S., Barneche, F., Peltier, G., and Rochaix, J. (2005). State transitions and light adaptation require chloroplast thylakoid protein kinase STN7. Nature 433, 892–895.

Bilger, W. and Björkman, O. (1990). Role of the xanthophyll cycle in photoprotection elucidated by measurements of light-induced absorbance changes, fluorescence and photosynthesis in leaves of Hedera canariensis. Photosynth Res. 25, 173–185.

Breyton, C., Nandha, B., Johnson, G.N., Joliot, P., and Finazzi, G. (2006). Redox modulation of cyclic electron flow around photosystem I in C3 plants. Biochemistry (N. Y.) 45, 13465–13475.

Brooks, M.D., Sylak-Glassman, E.J., Fleming, G.R., and Niyogi, K.K. (2013). A thioredoxin-like/beta-propeller protein maintains the efficiency of light harvesting in Arabidopsis. Proc. Natl. Acad. Sci. U. S. A. 110, 2733–2740.

Buchanan, B.B. (2016). The Path to Thioredoxin and Redox Regulation in Chloroplasts. 67, 1–24.

Carrillo, L.R., Froehlich, J.E., Cruz, J.A., Savage, L.J., and Kramer, D.M. (2016). Multi-level regulation of the chloroplast ATP synthase: the chloroplast NADPH thioredoxin reductase C (NTRC) is required for redox modulation specifically under low irradiance. Plant J. 87, 654–663.

Collin, V., Issakidis-Bourguet, E., Marchand, C., Hirasawa, M., Lancelin, J., Knaff, D., and Miginiac-Maslow, M. (2003). The Arabidopsis plastidial thioredoxins - New functions and new insights into specificity. J. Biol. Chem. 278, 23747–23752.

Courteille, A., Vesa, S., Sanz-Barrio, R., Cazale, A., Becuwe-Linka, N., Farran, I., Havaux, M., Rey, P., and Rumeau, D. (2013). Thioredoxin m4 Controls Photosynthetic Alternative Electron Pathways in Arabidopsis. Plant Physiol. 161, 508–520.

Couturier, J., Chibani, K., Jacquot, J., and Rouhier, N. (2013). Cysteine-based redox regulation and signaling in plants. Front. Plant Sci. 4, 105.

Cruz, J., Sacksteder, C., Kanazawa, A., and Kramer, D. (2001). Contribution of electric field (Delta psi) to steady-state transthylakoid proton motive force (pmf) in vitro and in vivo. Control of pmf parsing into Delta psi and Delta pH by ionic strength. Biochemistry (N. Y.) 40, 1226–1237.

Cruz, J., Avenson, T., Kanazawa, A., Takizawa, K., Edwards, G., and Kramer, D. (2005). Plasticity in light reactions of photosynthesis for energy production and photoprotection. J. Exp. Bot. 56, 395–406.

Da, Q., Sun, T., Wang, M., Jin, H., Li, M., Feng, D., Wang, J., Wang, H., and Liu, B. (2017). M-type thioredoxins are involved in the xanthophyll cycle and proton motive force to alter NPQ under low-light conditions in Arabidopsis. Plant Cell Rep. 37, 279–291.

DalCorso, G., Pesaresi, P., Masiero, S., Aseeva, E., Schuenemann, D., Finazzi, G., Joliot, P., Barbato, R., and Leister, D. (2008). A complex containing PGRL1 and PGR5 is involved in the switch between linear and cyclic electron flow in Arabidopsis. Cell 132, 273–285.

Demmig-Adams, B., Cohu, C.M., Muller, O., and Adams, W.W., 3rd (2012). Modulation of photosynthetic energy conversion efficiency in nature: from seconds to seasons. Photosynth Res. 113, 75–88.

Fan, D., Nie, Q., Hope, A.B., Hillier, W., Pogson, B.J., and Chow, W.S. (2007). Quantification of cyclic electron flow around Photosystem I in spinach leaves during photosynthetic induction. Photosynthesis Res. 94, 347–357.

Geigenberger, P., Thormählen, I., Daloso, D.M., and Fernie, A.R. (2017). The Unprecedented Versatility of the Plant Thioredoxin System. Trends Plant Sci. 22, 249–262.

Gollan, P.J., Lima-Melo, Y., Tiwari, A., Tikkanen, M., and Aro, E. (2017). Interaction between photosynthetic electron transport and chloroplast sinks triggers protection and signalling important for plant productivity. Philos Trans R Soc Lond B Biol Sci 372, 20160390, doi/10.1098/rstb.2016.0390.

Gotoh, E., Matsumoto, M., Ogawa, K., Kobayashi, Y., and Tsuyama, M. (2010). A qualitative analysis of the regulation of cyclic electron flow around photosystem I from the post-illumination chlorophyll fluorescence transient in Arabidopsis: a new platform for the in vivo investigation of the chloroplast redox state. Photosynthesis Res. 103, 111–123.

Grieco, M., Tikkanen, M., Paakkarinen, V., Kangasjärvi, S., and Aro, E.M. (2012). Steady-state phosphorylation of light-harvesting complex II proteins preserves photosystem I under fluctuating white light. Plant Physiol. 160, 1896–1910.

Hall, M., Mata-Cabana, A., Åkerlund, H.E., Florencio, F.J., Schroder, W.P., Lindahl, M., and Kieselbach, T. (2010). Thioredoxin targets of the plant chloroplast lumen and their implications for plastid function. Proteomics 10, 987–1001.

Hangarter, R.P., and Good, N.E. (1982) Energy thresholds for ATP synthesis in chloroplasts. Biochim. Biophys. Acta 681, 397–404

Hasan, S.S., Yamashita, E., Baniulis, D., and Cramer, W.A. (2013) Quinone-dependent proton transfer pathways in the photosynthetic cytochrome b_6_f complex. Proc. Natl. Acad. Sci. U. S. A. 110: 4297–4302.

Hertle, A.P., Blunder, T., Wunder, T., Pesaresi, P., Pribil, M., Armbruster, U., and Leister, D. (2013). PGRL1 Is the Elusive Ferredoxin-Plastoquinone Reductase in Photosynthetic Cyclic Electron Flow. Mol. Cell 49, 511–523.

Hirasawa, M., Schurmann, P., Jacquot, J., Manieri, W., Jacquot, P., Keryer, E., Hartman, F., and Knaff, D. (1999). Oxidation-reduction properties of chloroplast thioredoxins, Ferredoxin : Thioredoxin reductase, and thioredoxin f-regulated enzymes. Biochemistry (N. Y.) 38, 5200–5205.

Horvath, E., Peter, S., Joet, T., Rumeau, D., Cournac, L., Horvath, G., Kavanagh, T., Schafer, C., Peltier, G., and Medgyesy, P. (2000). Targeted inactivation of the plastid ndhB gene in tobacco results in an enhanced sensitivity of photosynthesis to moderate stomatal closure. Plant Physiol. 123, 1337–1349.

Johnson, G.N. (2011). Physiology of PSI cyclic electron transport in higher plants. Biochim. Biophys. Acta 1807, 384–389.

Joliot, P. and Joliot, A. (2002). Cyclic electron transfer in plant leaf. Proc. Natl. Acad. Sci. U. S. A. 99, 10209–10214.

Joliot, P. and Johnson, G.N. (2011). Regulation of cyclic and linear electron flow in higher plants. Proc. Natl. Acad. Sci. U. S. A. 108, 13317–13322.

Kanazawa, A., Ostendorf, E., Kohzuma, K., Hoh, D., Strand, D.D., Sato-Cruz, M., Savage, L., Cruz, J.A., Fisher, N., Froehlich, J.E., and Kramer, D.M. (2017). Chloroplast ATP Synthase Modulation of the Thylakoid Proton Motive Force: Implications for Photosystem I and Photosystem II Photoprotection. Front. Plant Sci. 8, 719.

Kirchsteiger, K., Pulido, P., Gonzalez, M., and Javier Cejudo, F. (2009). NADPH Thioredoxin Reductase C Controls the Redox Status of Chloroplast 2-Cys Peroxiredoxins in Arabidopsis thaliana. Mol. Plant. 2, 298–307.

Klughammer, C., and Schreiber, U. (2008a) Complementary PSII quantum yields calculated from simple fluorescence parameters measured by PAM fluorometry and the Saturation Pulse method. PAM Application Notes 1: 27–35.

Klughammer, C., and Schreiber, U. (2008b) Saturation Pulse method for assessment of energy conversion in PSI. PAM Application Notes 1: 11–14.

Klughammer, C., and Schreiber, U. (2016) Deconvolution of ferredoxin, plastocyanin, and P700 transmittance changes in intact leaves with a new type of kinetic LED array spectrophotometer. Photosynthesis. Res. 128: 195–214.

Klughammer, C., Siebke, K., and Schreiber, U. (2013). Continuous ECS-indicated recording of the proton-motive charge flux in leaves. Photosynthesis Res. 117, 471–487.

Kono, M. and Terashima, I. (2014). Long-term and short-term responses of the photosynthetic electron transport to fluctuating light. J. Photochem. Photobiol. B. 137, 89–99.

Kou, J., Takahashi, S., Fan, D., Badger, M.R., and Chow, W.S. (2015). Partially dissecting the steady-state electron fluxes in Photosystem I in wild-type and pgr5 and ndh mutants of Arabidopsis. Front. Plant Sci. 6, 758.

Kramer, D.M. and Crofts, A.R. (1989). Activation of the Chloroplast Atpase Measured by the Electrochromic Change in Leaves of Intact Plants. Biochim. Biophys. Acta 976, 28–41.

Kramer, D.M., Johnson, G., Kiirats, O., and Edwards, G.E. (2004). New fluorescence parameters for the determination of Q(A) redox state and excitation energy fluxes. Photosynthesis Res. 79, 209–218.

Kramer, D., Wise, R., Frederick, J., Alm, D., Hesketh, J., Ort, D., and Crofts, A. (1990). Regulation of Coupling Factor in Field-Grown Sunflower - a Redox Model Relating Coupling Factor Activity to the Activities of Other Thioredoxin-Dependent Chloroplast Enzymes. Photosynthesis Res. 26, 213–222.

Leister, D. and Shikanai, T. (2013). Complexities and protein complexes in the antimycin A-sensitive pathway of cyclic electron flow in plants. Front. Plant Sci. 4, 161.

Lepistö, A., Kangasjärvi, S., Luomala, E., Brader, G., Sipari, N., Keränen, M., Keinänen, M., and Rintamäki, E. (2009). Chloroplast NADPH-Thioredoxin Reductase Interacts with Photoperiodic Development in Arabidopsis. Plant Physiol. 149, 1261–1276.

Lepistö, A., Pakula, E., Toivola, J., Krieger-Liszkay, A., Vignols, F., and Rintamäki, E. (2013). Deletion of chloroplast NADPH-dependent thioredoxin reductase results in inability to regulate starch synthesis and causes stunted growth under short-day photoperiods. J. Exp. Bot. 64, 3843–3854.

Martin, M., Noarbe, D.M., Serrot, P.H., and Sabater, B. (2015). The rise of the photosynthetic rate when light intensity increases is delayed in ndh gene-defective tobacco at high but not at low CO2 concentrations. Front. Plant Sci. 6, 34.

Martinsuo, P., Pursiheimo, S., Aro, E., and Rintamaki, E. (2003). Dithiol oxidant and disulfide reductant dynamically regulate the phosphorylation of light-harvesting complex II proteins in thylakoid membranes. Plant Physiol. 133, 37–46.

McKinney, D., Buchanan, B., and Wolosiuk, R. (1978). Activation of Chloroplast Atpase by Reduced Thioredoxin. Phytochemistry 17, 794–795.

Miyake, C., Shinzaki, Y., Miyata, M., and Tomizawa, K. (2004). Enhancement of cyclic electron flow around PSI at high light and its contribution to the induction of non-photochemical quenching of chl fluorescence in intact leaves of tobacco plants. Plant Cell Physiol 45, 1426–1433.

Munekage, Y., Hojo, M., Meurer, J., Endo, T., Tasaka, M., and Shikanai, T. (2002). PGR5 is involved in cyclic electron flow around photosystem I and is essential for photoprotection in Arabidopsis. Cell 110, 361–371.

Munekage, Y., Hashimoto, M., Miyaka, C., Tomizawa, K., Endo, T., Tasaka, M., and Shikanai, T. (2004). Cyclic electron flow around photosystem I is essential for photosynthesis. Nature 429, 579–582.

Murashige, T. and Skoog, F. (1962). A Revised Medium for Rapid Growth and Bio Assays with Tobacco Tissue Cultures. Physiol. Plantarum 15, 473–497.

Nalin, C. and McCarty, R. (1984). Role of a Disulfide Bond in the Gamma-Subunit in Activation of the Atpase of Chloroplast Coupling Factor-i. J. Biol. Chem. 259, 7275–7280.

Naranjo, B., Mignee, C., Krieger-Liszkay, A., Hornero-Mendez, D., Gallardo-Guerrero, L., Javier Cejudo, F., and Lindahl, M. (2016). The chloroplast NADPH thioredoxin reductase C, NTRC, controls non-photochemical quenching of light energy and photosynthetic electron transport in Arabidopsis. Plant Cell Environ. 39, 804–822.

Nikkanen, L., Toivola, J., and Rintamäki, E. (2016). Crosstalk between chloroplast thioredoxin systems in regulation of photosynthesis. Plant Cell Environ. 39, 1691–1705.

Niyogi, K.K. and Truong, T.B. (2013). Evolution of flexible non-photochemical quenching mechanisms that regulate light harvesting in oxygenic photosynthesis. Curr. Opin. Plant Biol. 16, 307–314.

Okegawa, Y., Kagawa, Y., Kobayashi, Y., and Shikanai, T. (2008). Characterization of factors affecting the activity of photosystem I cyclic electron transport in chloroplasts. Plant Cell Physiol 825–834.

Peltier, G., Aro, E., and Shikanai, T. (2016). NDH-1 and NDH-2 Plastoquinone Reductases in Oxygenic Photosynthesis. Annual Review of Plant Biology, Annu. Rev. Plant Biol. 67, 55–80.

Perez-Ruiz, J.M., Spinola, M.C., Kirchsteiger, K., Moreno, J., Sahrawy, M., and Cejudo, F.J. (2006). Rice NTRC is a high-efficiency redox system for chloroplast protection against oxidative damage. Plant Cell 18, 2356–2368.

Pérez-Ruiz, J.M., Naranjo, B., Ojeda, V., Guinea, M., and Cejudo, F.J. (2017). NTRC-dependent redox balance of 2-Cys peroxiredoxins is needed for optimal function of the photosynthetic apparatus. Proc. Natl. Acad. Sci. U. S. A. 114, 12069–12074.

Petroutsos, D., Terauchi, A.M., Busch, A., Hirschmann, I., Merchant, S.S., Finazzi, G., and Hippler, M. (2009). PGRL1 Participates in Iron-induced Remodeling of the Photosynthetic Apparatus and in Energy Metabolism in Chlamydomonas reinhardtii. J. Biol. Chem. 284, 32770–32781.

Porra, R., Thompson, W., and Kriedemann, P. (1989). Determination of Accurate Extinction Coefficients and Simultaneous-Equations for Assaying Chlorophyll-a and Chlorophyll-B Extracted with 4 Different Solvents - Verification of the Concentration of Chlorophyll Standards by Atomic-Absorption Spectroscopy. Biochim. Biophys. Acta 975, 384–394.

Pulido, P., Spinola, M.C., Kirchsteiger, K., Guinea, M., Belen Pascual, M., Sahrawy, M., Sandalio, L.M., Dietz, K., Gonzalez, M., and Cejudo, F.J. (2010). Functional analysis of the pathways for 2-Cys peroxiredoxin reduction in Arabidopsis thaliana chloroplasts. J. Exp. Bot. 61, 4043–4054.

Richter, A.S., Peter, E., Rothbart, M., Schlicke, H., Toivola, J., Rintamäki, E., and Grimm, B. (2013). Posttranslational Influence of NADPH-Dependent Thioredoxin Reductase C on Enzymes in Tetrapyrrole Synthesis. Plant Physiol. 162, 63–73.

Rintamäki, E., Salonen, M., Suoranta, U., Carlberg, I., Andersson, B., and Aro, E. (1997). Phosphorylation of light-harvesting complex II and photosystem II core proteins shows different irradiance-dependent regulation in vivo - Application of phosphothreonine antibodies to analysis of thylakoid phosphoproteins. J. Biol. Chem. 272, 30476–30482.

Rintamäki, E., Martinsuo, P., Pursiheimo, S., and Aro, E. (2000). Cooperative regulation of light-harvesting complex II phosphorylation via the plastoquinol and ferredoxin-thioredoxin system in chloroplasts. Proc. Natl. Acad. Sci. U. S. A. 97, 11644–11649.

Rochaix, J. (2011). Regulation of photosynthetic electron transport. Biochim. Biophys. Acta-Bioenerg. 1807, 375–383.

Ruban, A.V. (2016). Nonphotochemical Chlorophyll Fluorescence Quenching: Mechanism and Effectiveness in Protecting Plants from Photodamage. Plant Physiol. 170, 1903–1916.

Ruban, A.V. and Johnson, M.P. (2009). Dynamics of higher plant photosystem cross-section associated with state transitions. Photosynthesis Res. 99, 173–183.

Rumeau, D., Becuwe-Linka, N., Beyly, A., Louwagie, M., Garin, J., and Peltier, G. (2005). New subunits NDH-M, -N, and -O, encoded by nuclear genes, are essential for plastid Ndh complex functioning in higher plants. Plant Cell 17, 219–232.

Schansker, G. and Strasser, R.J. (2005) Quantificaiton of non-Q_B_-reducing centers in leaves using farred pre-illumination. Photosynthesis Res. 84: 145–151.

Schneider, C.A., Rasband, W.S., and Eliceiri, K.W. (2012). NIH Image to ImageJ: 25 years of image analysis. Nat. Methods 9, 671–675.

Schreiber, U., and Klughammer, C. (2016) Analysis of Photosystem I donor and acceptor sides with a new type of online-deconvoluting kinetic LED-array spectrophotometer. Plant Cell Physiol. 57: 1454–1467.

Schreiber, U. (2017). Redox changes of ferredoxin, P700 and plastocyanin measured simultaneously in intact leaves. Photosynthesis Res. 134: 343–360.

Schreiber, U. and Klughammer, C. (2008). New accessory for the DUAL-PAM-100: The P515/535 module and examples of its application. PAM Application Notes 1, 1–10.

Schürmann, P. and Buchanan, B.B. (2008). The ferredoxin/thioredoxin system of oxygenic photosynthesis. Antioxid. Redox Signal. 10, 1235–1273.

Serrato, A.J., Perez-Ruiz, J.M., Spinola, M.C., and Cejudo, F.J. (2004). A novel NADPH thioredoxin reductase, localized in the chloroplast, which deficiency causes hypersensitivity to abiotic stress in Arabidopsis thaliana. J. Biol. Chem. 279, 43821–43827.

Shapiguzov, A., Chai, X., Fucile, G., Longoni, P., Zhang, L., and Rochaix, J.-. (2016). Activation of the Stt7/STN7 Kinase through Dynamic Interactions with the Cytochrome b(6)f Complex. Plant Physiol. 171, 1533–1533.

Shikanai, T., Endo, T., Hashimoto, T., Yamada, Y., Asada, K., and Yokota, A. (1998). Directed disruption of the tobacco ndhB gene impairs cyclic electron flow around photosystem I. Proc. Natl. Acad. Sci. U. S. A. 95, 9705–9709.

Shikanai, T. (2016). Chloroplast NDH: A different enzyme with a structure similar to that of respiratory NADH dehydrogenase. Biochim. Biophys. Acta-Bioenerg. 1857, 1015–1022.

Shikanai, T. and Yamamoto, H. (2017). Contribution of Cyclic and Pseudo-cyclic Electron Transport to the Formation of Proton Motive Force in Chloroplasts. Mol. Plant. 10, 20–29.

Sievers, F., Wilm, A., Dineen, D., Gibson, T.J., Karplus, K., Li, W., Lopez, R., McWilliam, H., Remmert, M., Soeding, J., Thompson, J.D., and Higgins, D.G. (2011). Fast, scalable generation of high-quality protein multiple sequence alignments using Clustal Omega. Mol. Syst. Biol. 7, 539.

Stirbet, A., Riznichenko, G.Y., Rubin, A.B., and Govindjee (2014). Modeling chlorophyll a fluorescence transient: relation to photosynthesis. Biochemistry (Mosc.) 79, 291–323.

Strand, D.D., Fisher, N., and Kramer, D.M. (2017a). The higher plant plastid NAD(P)H dehydrogenase-like complex (NDH) is a high efficiency proton pump that increases ATP production by cyclic electron flow. J. Biol. Chem. 292, 11850–11860.

Strand, D.D., Livingston, A.K., Satoh-Cruz, M., Froehlich, J.E., Maurino, V.G., and Kramer, D.M. (2015). Activation of cyclic electron flow by hydrogen peroxide in vivo. Proc. Natl. Acad. Sci. U. S. A. 112, 5539–5544.

Strand, D.D., Fisher, N., Davis, G.A., and Kramer, D.M. (2016a). Redox regulation of the antimycin A sensitive pathway of cyclic electron flow around photosystem I in higher plant thylakoids. Biochim. Biophys. Acta-Bioenerg. 1857, 1–6.

Strand, D.D., Fisher, N., and Kramer, D.M. (2016b). Distinct Energetics and Regulatory Functions of the Two Major Cyclic Electron Flow Pathways in Chloroplasts In Chloroplasts: Current Research and Future Trends, Kirchhoff, H., ed (Wymondham; 32 Hewitts Lane, Wymondham Nr 18 0JA, England: Caister Academic Press) pp. 89–100.

Strand, D.D., Livingston, A.K., Satoh-Cruz, M., Koepke, T., Enlow, H.M., Fisher, N., Froehlich, J.E., Cruz, J.A., Minhas, D., Hixson, K.K., Kohzuma, K., Lipton, M., Dhingra, A., and Kramer, D.M. (2017b). Defects in the Expression of Chloroplast Proteins Leads to H2O2 Accumulation and Activation of Cyclic Electron Flow around Photosystem I. Front. Plant Sci. 7, 2073.

Suorsa, M., Jarvi, S., Grieco, M., Nurmi, M., Pietrzykowska, M., Rantala, M., Kangasjarvi, S., Paakkarinen, V., Tikkanen, M., Jansson, S., and Aro, E. (2012). PROTON GRADIENT REGULATION5 Is Essential for Proper Acclimation of Arabidopsis Photosystem I to Naturally and Artificially Fluctuating Light Conditions. Plant Cell 24, 2934–2948.

Suorsa, M., Grieco, M., Järvi, S., Gollan, P.J., Kangasjärvi, S., Tikkanen, M., and Aro, E. (2013). PGR5 ensures photosynthetic control to safeguard photosystem I under fluctuating light conditions. Plant Signal. Behav. 8, e22714.

Suorsa, M. (2015). Cyclic electron flow provides acclimatory plasticity for the photosynthetic machinery under various environmental conditions and developmental stages. Front. Plant Sci. 6, 800.

Suorsa, M., Rossi, F., Tadini, L., Labs, M., Colombo, M., Jahns, P., Kater, M.M., Leister, D., Finazzi, G., Aro, E., Barbato, R., and Pesaresi, P. (2016). PGR5-PGRL1-Dependent Cyclic Electron Transport Modulates Linear Electron Transport Rate in Arabidopsis thaliana. Mol. Plant. 9, 271–288.

Takagi, D., and Myake, C. (2018) Proton gradient regulation 5 supports linear electron flow to oxidize photosystem I. Physiol. Plantarum, in press, doi: 10.1111/ppl.12723.

Thormählen, I., Meitzel, T., Groysman, J., Ochsner, A.B., von Roepenack-Lahaye, E., Naranjo, B., Cejudo, F.J., and Geigenberger, P. (2015). Thioredoxin f1 and NADPH-dependent thioredoxin reductase C have overlapping functions in regulating photosynthetic metabolism and plant growth in response to varying light conditions. Plant Physiol. 169, 1766–1786.

Thormählen, I., Zupok, A., Rescher, J., Leger, J., Weissenberger, S., Groysman, J., Orwat, A., Chatel-Innocenti, G., Issakidis-Bourguet, E., Armbruster, U., and Geigenberger, P. (2017). Thioredoxins Play a Crucial Role in Dynamic Acclimation of Photosynthesis in Fluctuating Light. Mol. Plant. 10, 168–182.

Tikkanen, M. and Aro, E.M. (2014). Integrative regulatory network of plant thylakoid energy transduction. Trends Plant Sci. 19, 10–17.

Tikkanen, M., Pippo, M., Suorsa, M., Sirpio, S., Mulo, P., Vainonen, J., Vener, A., Allahverdiyeva, Y., and Aro, E.M. (2006). State transitions revisited - a buffering system for dynamic low light acclimation of Arabidopsis. Plant Mol. Biol. 62, 779–793.

Tikkanen, M., Grieco, M., Kangasjarvi, S., and Aro, E. (2010). Thylakoid Protein Phosphorylation in Higher Plant Chloroplasts Optimizes Electron Transfer under Fluctuating Light. Plant Physiol. 152, 723–735.

Tikkanen, M., Grieco, M., Nurmi, M., Rantala, M., Suorsa, M., and Aro, E. (2012). Regulation of the photosynthetic apparatus under fluctuating growth light. Philos. Trans. R. Soc. Lond. B Biol. Sci. 367, 3486–3493.

Tikkanen, M., Rantala, S., and Aro, E. (2015). Electron flow from PSII to PSI under high light is controlled by PGR5 but not by PSBS. Front. Plant Sci. 6, 521.

Tiwari, A., Mamedov, F., Grieco, M., Suorsa, M., Jajoo, A., Styring, S., Tikkanen, M., and Aro, E. (2016). Photodamage of iron-sulphur clusters in photosystem I induces non-photochemical energy dissipation. Nat. Plants 2, 16035.

Toivola, J., Nikkanen, L., Dahlström, K.M., Salminen, T.A., Lepistö, A., Vignols, F., and Rintamäki, E. (2013). Overexpression of chloroplast NADPH-dependent thioredoxin reductase in Arabidopsis enhances leaf growth and elucidates in vivo function of reductase and thioredoxin domains. Front. Plant Sci. 4, 389.

Toth, S.Z., Schansker, G., and Strasser, R.J. (2007). A non-invasive assay of the plastoquinone pool redox state based on the OJIP-transient. Photosynthesis Res. 93, 193–203.

Townsend, A.J., Ware, M.A., and Ruban, A.V. (2017). Dynamic interplay between photodamage and photoprotection in photosystem II. Plant. Cell. Environ. 41, 1098–1112

Trotta, A., Suorsa, M., Rantala, M., Lundin, B., and Aro, E. (2016). Serine and threonine residues of plant STN7 kinase are differentially phosphorylated upon changing light conditions and specifically influence the activity and stability of the kinase. Plant J. 87, 484–494.

Vener, A., VanKan, P., Rich, P., Ohad, I., and Andersson, B. (1997). Plastoquinol at the quinol oxidation site of reduced cytochrome bf mediates signal transduction between light and protein phosphorylation: Thylakoid protein kinase deactivation by a single-turnover flash. Proc. Natl. Acad. Sci. U. S. A. 94, 1585–1590.

Walter, M., Chaban, C., Schutze, K., Batistic, O., Weckermann, K., Nake, C., Blazevic, D., Grefen, C., Schumacher, K., Oecking, C., Harter, K., and Kudla, J. (2004). Visualization of protein interactions in living plant cells using bimolecular fluorescence complementation. Plant J. 40, 428–438.

Wang, C., Yamamoto, H., and Shikanai, T. (2015). Role of cyclic electron transport around photosystem I in regulating proton motive force. Biochim. Biophys. Acta-Bioenerg. 1847, 931–938.

Wang, P., Liu, J., Liu, B., Da, Q., Feng, D., Su, J., Zhang, Y., Wang, J., and Wang, H. (2014). Ferredoxin: Thioredoxin Reductase Is Required for Proper Chloroplast Development and Is Involved in the Regulation of Plastid Gene Expression in Arabidopsis thaliana. Mol. Plant. 7, 1586–1590.

Yamamoto, H., Peng, L., Fukao, Y., and Shikanai, T. (2011). An Src Homology 3 Domain-Like Fold Protein Forms a Ferredoxin Binding Site for the Chloroplast NADH Dehydrogenase-Like Complex in Arabidopsis. Plant Cell 23, 1480–1493.

Yamamoto, H. and Shikanai, T. (2013). In Planta Mutagenesis of Src Homology 3 Domain-like Fold of NdhS, a Ferredoxin-binding Subunit of the Chloroplast NADH Dehydrogenase-like Complex in Arabidopsis A conserved Arg-193 playsa critical role in ferredoxin binding. J. Biol. Chem. 288, 36328–36337.

Yamori, W., Sakata, N., Suzuki, Y., Shikanai, T., and Makino, A. (2011). Cyclic electron flow around photosystem I via chloroplast NAD(P)H dehydrogenase (NDH) complex performs a significant physiological role during photosynthesis and plant growth at low temperature in rice. Plant J. 68, 966–976.

Yamori, W., Shikanai, T., and Makino, A. (2015). Photosystem I cyclic electron flow via chloroplast NADH dehydrogenase-like complex performs a physiological role for photosynthesis at low light. Sci. Rep. 5, 15593.

Yamori, W., Makino, A., and Shikanai, T. (2016). A physiological role of cyclic electron transport around photosystem I in sustaining photosynthesis under fluctuating light in rice. Sci. Rep. 6, 20147.

Yamori, W. and Shikanai, T. (2016). Physiological Functions of Cyclic Electron Transport Around Photosystem I in Sustaining Photosynthesis and Plant Growth. Annu. Rev. Plant Biol. 67, 81–106.

Yoshida, K., Hara, S., and Hisabori, T. (2015). Thioredoxin Selectivity for Thiol-Based Redox Regulation of Target Proteins in Chloroplasts. J. Biol. Chem. 290, 19540.

Yoshida, K. and Hisabori, T. (2016). Two distinct redox cascades cooperatively regulate chloroplast functions and sustain plant viability. Proc. Natl. Acad. Sci. U. S. A. 113, 3967–3976.

Yoshida, K. and Hisabori, T. (2017). Distinct Electron Transfer from Ferredoxin-Thioredoxin Reductase to Multiple Thioredoxin Isoforms in Chloroplasts. Biochem. J. 474, 1347–1360, doi/10.1042/BCJ20161089.

Zaffagnini, M., De Mia, M., Morisse, S., Di Giacinto, N., Marchand, C.H., Maes, A., Lemaire, S.D., and Trost, P. (2016). Protein S-nitrosylation in photosynthetic organisms: A comprehensive overview with future perspectives. Biochim. Biophys. Acta-Proteom. 1864, 952–966.

